# Sex chromosome identification and genome curation from a single individual with SCINKD

**DOI:** 10.1101/2025.07.07.660342

**Authors:** Brendan J. Pinto, Simone M. Gable, Shannon E. Keating, Chase H. Smith, Tony Gamble, Stuart V. Nielsen, Melissa A. Wilson

## Abstract

In most animal species, the sex determining pathway is typically initiated by the presence/absence of a primary genetic cue at a critical point during development. This primary genetic cue is often located on a single locus—referred to as sex chromosomes—and can be limited to females (in a ZZ/ZW system) or males (in an XX/XY system). One trademark of sex chromosomes is a restriction or cessation of recombination surrounding the sex-limited region (to prevent its inheritance in the homogametic sex). This may lead to—through a variety of mechanisms—higher amounts of genetic divergence within this region, i.e. between the X/Z and Y/W chromosomes, especially when compared to their autosomal counterparts. Recent advances in genome sequencing and computation have brought with them the ability to resolve haplotypes within a diploid individual, permitting assembly of previously challenging genomic regions like sex chromosomes. Leveraging these advances, we identified replicable diagnostic characteristics between typical autosomes and sex chromosomes (within a single genome assembly). Under this framework, we can use this information to identify putative sex chromosome linkage groups across divergent vertebrate taxa and simultaneously curate misassembled regions on autosomes. Here, we present this conceptual framework and associated tool for identifying candidate sex chromosome linkage groups from a single, diploid individual dubbed Sex Chromosome Identification by Negating Kmer Densities, or SCINKD.

## Introduction

In species that reproduce sexually, populations include individuals that produce eggs (females) or sperm (males). The process of sex determination can be initiated via environmental or genetic cues. In diploid vertebrates with environmental sex determination (ESD), all individuals possess two copies of each chromosome that can recombine over their entire length. In contrast, most vertebrate species possess sex chromosomes with genetic differences that lead to a developmental cascade becoming typical females and males, i.e. genetic sex determination. The locus containing a gene(s) of large effect in sex determination is often located in a region that has restricted recombination, or does not recombine, and can therefore be inherited either strictly maternally (W-specific in a ZZ/ZW system) or paternally (Y-specific in a XX/XY system). Over time, these sex-limited regions are often subjected to forces that suppress recombination and expand beyond this region where mutations accumulate quickly due to an inability to repair DNA damage through meiotic recombination. Indeed, we know that this mode of sex chromosome evolution is widespread in animals, if not the majority of systems (Graves, 2016; Jeffries et al., 2025; Marshall Graves, 2008).

Developing methods to quickly identify candidate sex-linked regions in genome assemblies at scale is of utmost importance as biodiversity genomics data across the tree of life are now being produced at unprecedented, albeit unequal, rates across the globe (Card et al., 2023; Gable et al., 2023; Linck & Cadena, 2024; Pinto, Gamble, Smith, & Wilson, 2023; Rhie et al., 2021).

Over the past ∼15 years, empiricists have developed many techniques to identify sex chromosomes using genetic sequence data from multiple female and male samples (Palmer et al., 2019). In the absence of diverse panels of reference-quality genomes for non-model organisms, the NGS-based revolution in sex chromosome identification began with RADseq (Feron et al., 2021; Gamble & Zarkower, 2014). Within the last decade, the progression towards whole-genome sequencing (WGS) data was accompanied by the requirement of a reference genome (Conte et al., 2017; Pinto et al., 2024). Most recently, the rise of the use of kmers in WGS data have pushed us back towards a reference-optional approach to identifying sex chromosomes (Behrens et al., 2024; Carey et al., 2024). However, with the advent of relatively inexpensive long read sequencing, generating diploid genome assemblies in biodiverse taxa has become more approachable, while the availability of population-level genomic sampling in many taxa has remained stagnant. Indeed, while often overlooked as sequencing has become easier and costs significantly decreased, field collections of population-level samples of plants, animals, and other eukaryotes have remained challenging. Thus, at its most extreme in this biodiversity genomics age, it is now possible to sequence and assemble the haplotype-resolved sex chromosome complement in a reference genome, but neglect to properly identify or annotate (formally label) them (see herein).

Prior DNA sequencing platforms (short-read and noisy long-read) were hindered by an inability to assemble high-quality genomes, especially the sex chromosomes, at scale. This issue has caused a historical, and intentional, bias towards sequencing mostly the homogametic sex (XX/ZZ individuals) in well-known taxa, such as mammals (XX), caenophidian snakes (ZZ), and birds (ZZ) to improve contiguity in *de novo* genome assemblies (Carvalho & Clark, 2013). However, cutting-edge sequencing technologies and computational methods have brought unprecedented insight into genome form and function (Antipov et al., 2024; Cheng et al., 2022), permitting access to previously uncharacterized genomic regions, including sex chromosomes at scale (Makova et al., 2023; Rhie et al., 2023). However, characterization of sex chromosomes in non-model systems remains challenging and, again, currently relies on generating additional data beyond a reference genome assembly, which is prohibitive in many instances (Pinto et al., 2022, 2024; Pinto, Gamble, Smith, & Wilson, 2023). To address the gap between reference genome availability and supporting data, we developed an approach to identify sex chromosomes from a single, diploid individual’s genome (Figure 1). This method is also useful for identifying finer-scale phasing and assembly issues. Leveraging a few modest assumptions regarding within-genome correlations, which appear to be conserved across most vertebrates, we describe Sex Chromosome Identification by Negating Kmer Densities (SCINKD) to identify unannotated sex chromosomes and curate diploid genome assemblies from a single individual.

**Figure 1:**
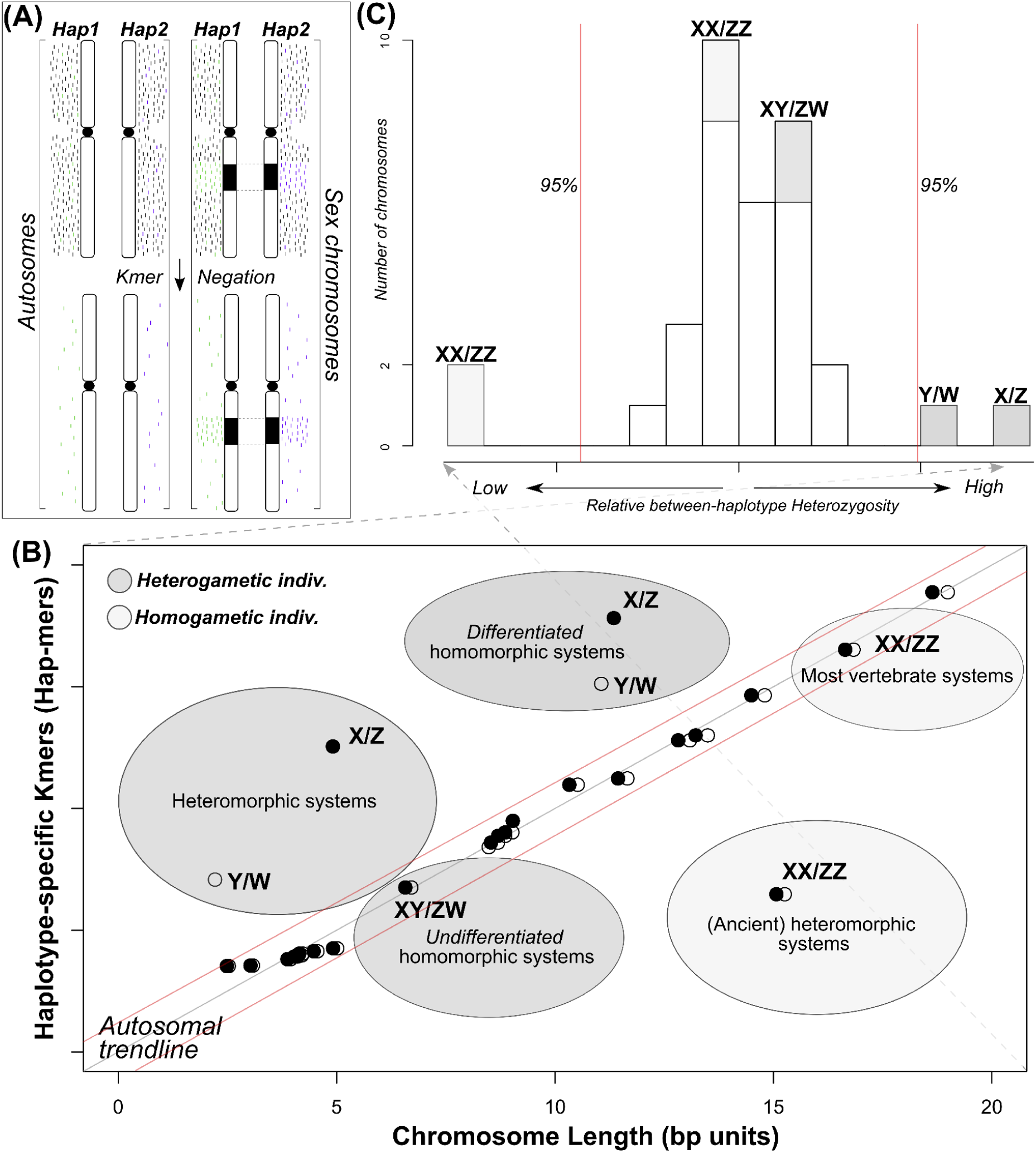
Simplified visual description of the SCINKD framework: (A) In many taxa, autosomes and sex chromosomes show disparate patterns where the region of restricted recombination (black) possesses increased haplotype-specific kmer (hap-mer) densities relative to the autosomal background. (B) This pattern drives overall increases in the number of hap-mers beyond what would be expected by chance (i.e. given the length of the chromosome). Each ‘bubble’ on the plot displays a hypothetical example of where the sex chromosomes of any given taxon could appear on the plot, including those that would be undetectable by SCINKD. (C) This deviation in hap-mers is detectable as a statistical outlier that can then be scrutinized using multivariate data visualization and/or additional data from samples of known sex.

## Methods

The SCINKD [v2.1] workflow is three steps, requiring as input a haplotype-resolved genome assembly using a haplotype-aware assembler, such as hifiasm or verkko (Antipov et al., 2024; Cheng et al., 2021, 2022), optimally scaffolded to chromosome-level. The steps are:

1. Count kmers (k=28) from both assembled haplotypes using meryl (Rhie et al., 2020)
2. Negate kmers that occur in both haplotype assemblies, keeping only kmers unique to each assembly.
3. Identify loci with kmer densities that deviate from the autosomal background using multivariate data visualization (dot plots, mirror plots, and haplotype alignments).

The tool to conduct these steps is developed as a snakemake workflow and template R files for data visualization; the FULL pipeline comes pre-optimized for most genomes (<∼5Gb) and is co-distributed with a GREEDY mode that uses homopolymer compression. However, the GREEDY mode remains largely untested for this purpose due to lack of high-quality assemblies of large genomes; it has potential uses with larger genomes (>5Gb). Both implementations come preconditioned for use on machines with 24 available threads and 24Gb of RAM. The full implementation of this workflow and a detailed tutorial on how to run and input requirements is available on GitHub (https://github.com/DrPintoThe2nd/SCINKD).

### Generation of new data

We extracted high molecular-weight DNA from the liver of adult, male *Sphaerodactylus notatus* via SOP-CEPC (Pinto et al., 2021) and generated a HiC library from the same individual using a DoveTail Omni-C kit (Cantata Bio; Cambridge, MA, USA). A single PacBio HiFi SMRTbell library was constructed from the extracted HMW DNA, which was subsequently barcoded and sequenced across 1.5 SMRT cells at the Arizona Genomics Institute (AGI) (University of Arizona; Tucson, AZ, USA). We sequenced the HiC library on an Illumina NovaSeq 6000 at the Texas A&M Agrilife Core Facility (College Station, TX, USA). We interrogated sequence features using fastqc (Andrews, 2010) and GenomeScope (Ranallo-Benavidez et al., 2020). All other data used in this study was published and/or downloaded from public access databases (Supplementary Table 1).

### Data analysis

Previous work showed extensive within-genome correlations using only a single reference genome (Pinto, Gamble, Smith, & Wilson, 2023). To corroborate whether a null expectation of the number of SNPs per chromosome as a function of chromosome length exists, we re-analyzed a small subset of human data. We downloaded data from Genotype-Tissue Expression (GTEx) project, approved for project #8834 for General Research Use in Genotype-Tissue Expression (GTEx) to MAW. We identified six individuals with an XX genotype in the GTEx data that matched specific search criteria for an independent project.

These data were archived in the genome-aligned CRAM format, thus, we re-processed them with a modified GRCh38 reference to account for sex chromosome complement following previously described methods (Pinto, O’Connor, et al., 2023) and calculated numbers of biallelic SNPs using rtg-tools (Cleary et al., 2015). We tested for significant differences between the number of SNPs across chr7 (159Mb) and chr8 (145Mb) versus chrX (156Mb) using the Wilcoxon rank-sum test (Wilcoxon, 1945).

We assembled HiFi reads with hifiasm [v0.24.0-r702] including HiC data for haplotype phasing (Cheng et al., 2021, 2022) for each test species: *Hemicordylus capensis* (Leitão et al., 2023), *Correlophus ciliatus* (Gumangan et al., 2024), *Eublepharis macularius* (Pinto, Gamble, Smith, Keating, et al., 2023)*, Lepidodactylus listeri* (Dodge et al., 2023), and *Sphaerodactylus notatus* (*this study*) (Supplemental Table 1). We scaffolded each haplotype separately using either juicebox [v1.6] (Durand et al., 2016) or Pretext (https://github.com/sanger-tol/PretextMap). We then conducted quality control of the assembly for gene content using BUSCO [v5.1.2] (Nishimura et al., 2017; Simão et al., 2015), calculated haplotype-specific statistics and visualizations by aligning the two haplotypes using minimap2 [v2.28-r1209] (Li, 2018) and SVbyEye (Porubsky et al., 2024). For *Podarcis cretensis* (ZZ), we reassembled the diploid genome using hifiasm and scaffolded by aligning to the NCBI reference genome using RagTag (Alonge et al., 2022).

For each species with individuals of known sex, we used SCINKD to identify putative sex chromosomes within each genome and cross-validated with either the literature, additional data, or both (Leitão et al., 2023; Pinto et al., 2022). To test whether or not SCINKD would work in a species without HiC data, we re-assembled the *Cryptoblepharus egeriae* genome, a species with higher quality data, but lacking HiC data (Dodge et al., 2023). Due to the limitations of assembling HiFi reads without HiC for phasing and scaffolding, we used the `--dual-scaf` setting in Hifiasm to generate comparable 1:1 contigs, we then filtered contigs less than 1Mb to reduce noise from genomic ‘shrapnel’. Lastly, for *Sphaerodactylus notatus* and *Correlophus ciliatus*, we leveraged recently developed whole-genome data with previously published gametolog-specific RADtags to supplement SCINKD results. To confirm the sex chromosome system identified via SCINKD, we mapped these RADtags (X and Y or Z and W, respectively) to each of their respective diploid assemblies (Gamble et al., 2015; Gumangan et al., 2024; Keating, 2022; Nielsen et al., 2019; Pinto et al., 2022).

To rigorously explore the sensitivity and features of SCINKD with regard to detecting misassemblies to flag for additional curation and the effects of sequencing depth and kmer length, we systematically interrogated the genome data of *Anniella stebbinsi* (a putative ZW female with a strong sex chromosome signal stemming from a ∼10Mb region at the distal end of chromosome 7) with two independent analyses. First, we re-assembled the haplotype-resolved reference genome using all available data (∼47x coverage) as detailed above and systematically hard-masked a region on chromosome 2 in haplotype 1 of varying sizes (1Mb, 5Mb, 10Mb, and 15Mb) and ran SCINKD on each of the four modified haplotype pairs (Supplemental Figure 1). Next, we returned to the raw PacBio data and subsampled it at varying coverages (20x, 25x, 30x, and 40x) using seqtk (https://github.com/lh3/seqtk). We assembled and scaffolded each of the eight haplotypes using HiFiasm and Pretext, respectively, as described previously. We then replicated SCINKD five times for each pair of haplotypes while varying kmer lengths: k21, k28 (default), k31, k41, and k51 (Supplemental Figure 2).

## Results and Discussion

Recent work examining within-genome correlations set strong expectations of linear relationships between chromosome length and content (GC content, gene density, SNPs, etc.) across vertebrates (Supplemental Figure 3) (Pinto, Gamble, Smith, & Wilson, 2023). In-line with this preliminary work, we hypothesized that this was also true for haplotype-specific kmers (hap-mers) in the leopard gecko (*Eublepharis macularius*), which is exactly what we found (R^2^ == 0.901) (Supplemental Figure 4). Importantly, the leopard gecko only possesses autosomes, as it is a species with a well-supported temperature dependent sex determination system (TSD). This finding gave rise to the prediction that hap-mers are correlated with chromosome length, except in regions that differ significantly in sequence composition, such as the non-recombining region(s) of the sex chromosomes or severely misassembled genomic regions (Figure 1A). To varying degrees, this is what we observe in our series of exemplar vertebrate datasets with known sex chromosome systems (Figure 2). Thus, hap-mer densities are strongly correlated between homologous autosomes, but can be significantly different between the sex chromosomes across many taxa (Figure 1).

**Figure 2:**
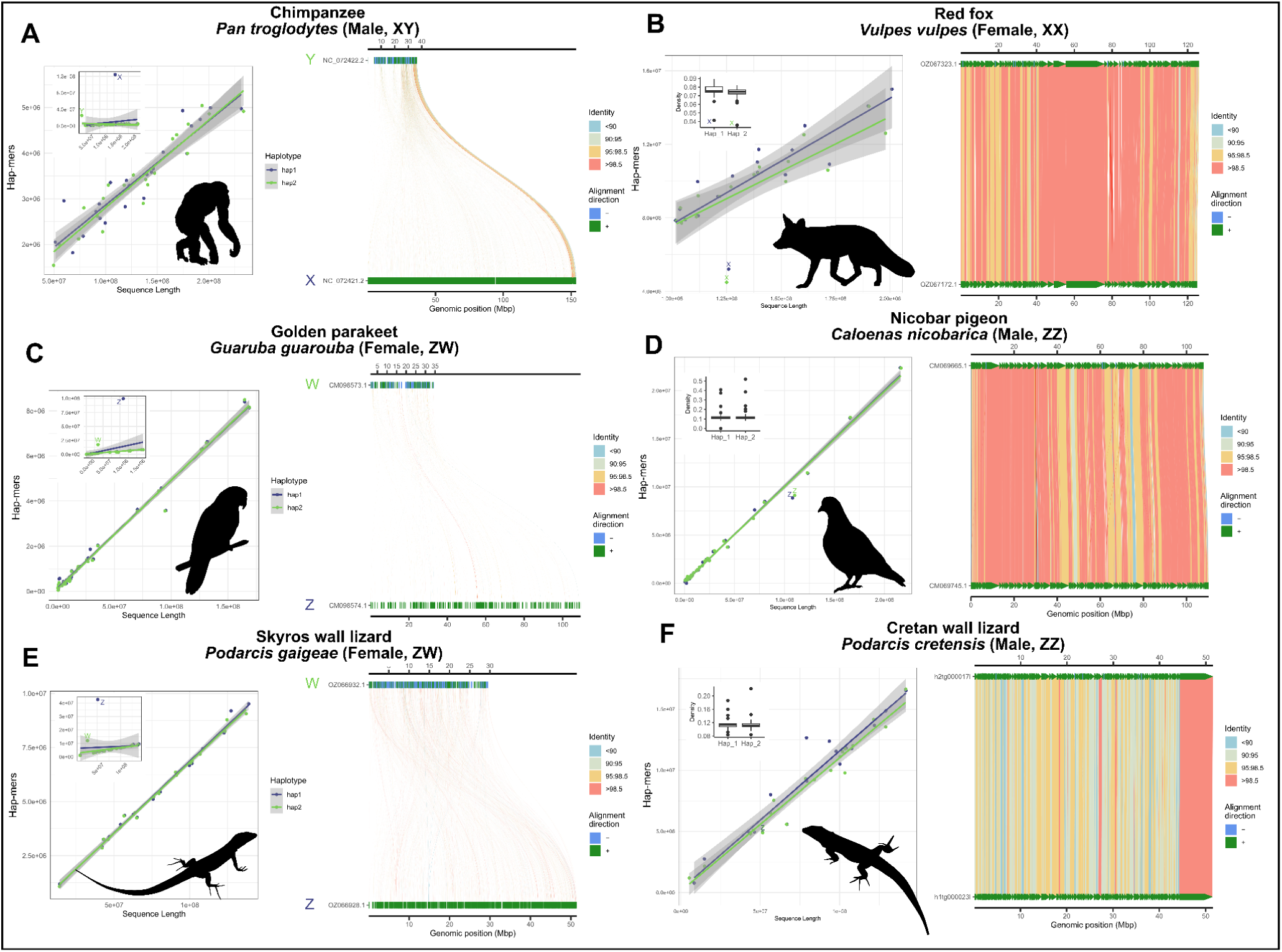
Exemplars of (A-B) mammal: XY (chimpanzee, *Pan troglodytes*) and XX (Red fox, *Vulpes vulpes*) individuals; (C-D) avian reptile: ZW (Golden parakeet, *Guaruba guarouba*) and ZZ (Nicobar pigeon, *Caloenas nicobarica*) individuals; (E-F) lacertid lizard: ZW (Skyros wall lizard, *Podarcis gaigeae*) and ZZ (Cretan wall lizard, *Podarcis cretensis*) individuals. Each panel consists of SCINKD “.results” output ∼ sequence length (Li et al., 2009) visualized using ggplot2 (Wickham, 2016) including only autosomes (exemplifying the predicted genome correlation), with an embedded panel demonstrating the full disparity between sex chromosomes and autosomes. Each panel also contains the corresponding minimap2 alignment (Li, 2018) of putative sex chromosome haplotypes visualized using SVbyEye (Porubsky et al., 2024).

### Previously known exemplar cases

We developed a series of test cases from well-known XY and ZW systems from each of the three major amniote lineages ordered by relative species number (low to high): mammals, birds, and squamates (Figure 2). Each has numerous high-quality genomic data, ranging from publicly available HiFi reads to complete T2T genomes. For each, we used the available GenBank assembly, or reassembled from raw data when haplotype-resolved information was unavailable. In mammals, we chose the well-characterized, telomere-to-telomere (T2T) chimpanzee (*Pan troglodytes*; XY) and the chromosome-level red fox (*Vulpes vulpes*; XX) genomes. In both species, sex chromosomes are apparent when examining dot plots of SCINKD results by chromosome length (Figure 2A-B). When aligning the two haplotypes for each species, we see the pseudoautosomal region (PAR) in chimpanzee aligns well, while the rest of the homologous pair aligns poorly. In the red fox, lower-than-expected heterozygosity on the pair of X chromosomes is exemplified by large stretches of high sequence similarity, replicating our expectations from human data (Supplemental Figure 3). In the two bird species, we see a similar pattern to chimpanzee in the Golden parakeet (ZW) and, to a much lesser extent, a similar pattern to the red fox in the Nicobar pigeon (ZZ) (Figure 2C-D). Both therian mammal and avian sex chromosome systems are ancient, heteromorphic systems that generally serve as exceptional, not typical, vertebrate sex chromosome systems.

To balance these extreme examples, we examined well-characterized systems in a more diverse group, squamate reptiles (Figure 2E-F). In lizards of the genus *Podarcis* (wall lizards), we see the signs of their heteromorphic ZZ/ZW system, which is conserved at the family level (Lacertidae) (Rovatsos et al., 2016). In the Skyros wall lizard (*P. gaigeae*; ZW), we observe a similar pattern to that of XY mammals and ZW birds, where the divergent nature of the heteromorphic system is apparent. However, in the Cretan wall lizard (*P. cretensis*; ZZ), the Z chromosomes show no visible diagnostic differences from any randomly selected pair of autosomes, which is a pattern expected for sex chromosome systems that are much more recently evolved than that of mammals or birds, and thus have not been subjected to the same kind of reduction in genetic diversity on the Z (Figure 1B) (Pinto et al., 2022). Lastly, we extended our inferences to a pair of much less divergent systems within the infraorder Scincomorpha. Here—unlike mammals, birds, and lacertids—both species have homomorphic sex chromosome systems, or systems that are indistinguishable under a light microscope. In the Cape cliff lizard (*Hemicordylus capensis*; XY), we see diagnostic differences between their X and Y chromosomes that are reinforced by multiple rearrangements between the chromosomal pair when aligned (Figure 3). And in the Christmas Island skink (*Cryptoblepharus egeriae*; XY), we observed a pair of contigs deviating from autosomal background on a region corresponding to the known sex chromosome linkage group in chicken (Gg1) (Figure 4). Embedded within these contigs is a small region that aligned poorly, the sex-limited region, corresponding to published expectations for the system in skinks (Kostmann et al., 2021). In passing, we note that within the publicly available contig assembly for the Christmas Island skink (GCA_030015325.1), there exists an equivalent contig to the chrX contig presented here, JAREYC010000008.1, while the putative sex-limited region of chrY is split off as its own unique contig, JAREYC010000009.1. Thus, both X/Z and Y/W gametologous regions possess detectably unique densities of hap-mers—not seen in their autosomal counterparts—in both highly degenerated heteromorphic systems that are readily diagnosable under a light microscope (e.g. mammals, birds, and lacertids; Figure 2), and those that aren’t, i.e. “homomorphic” systems (e.g. *Cryptoblepharus* and *Hemicordylus*; Figures 3 & 4) (Charlesworth, 2021).

**Figure 3:**
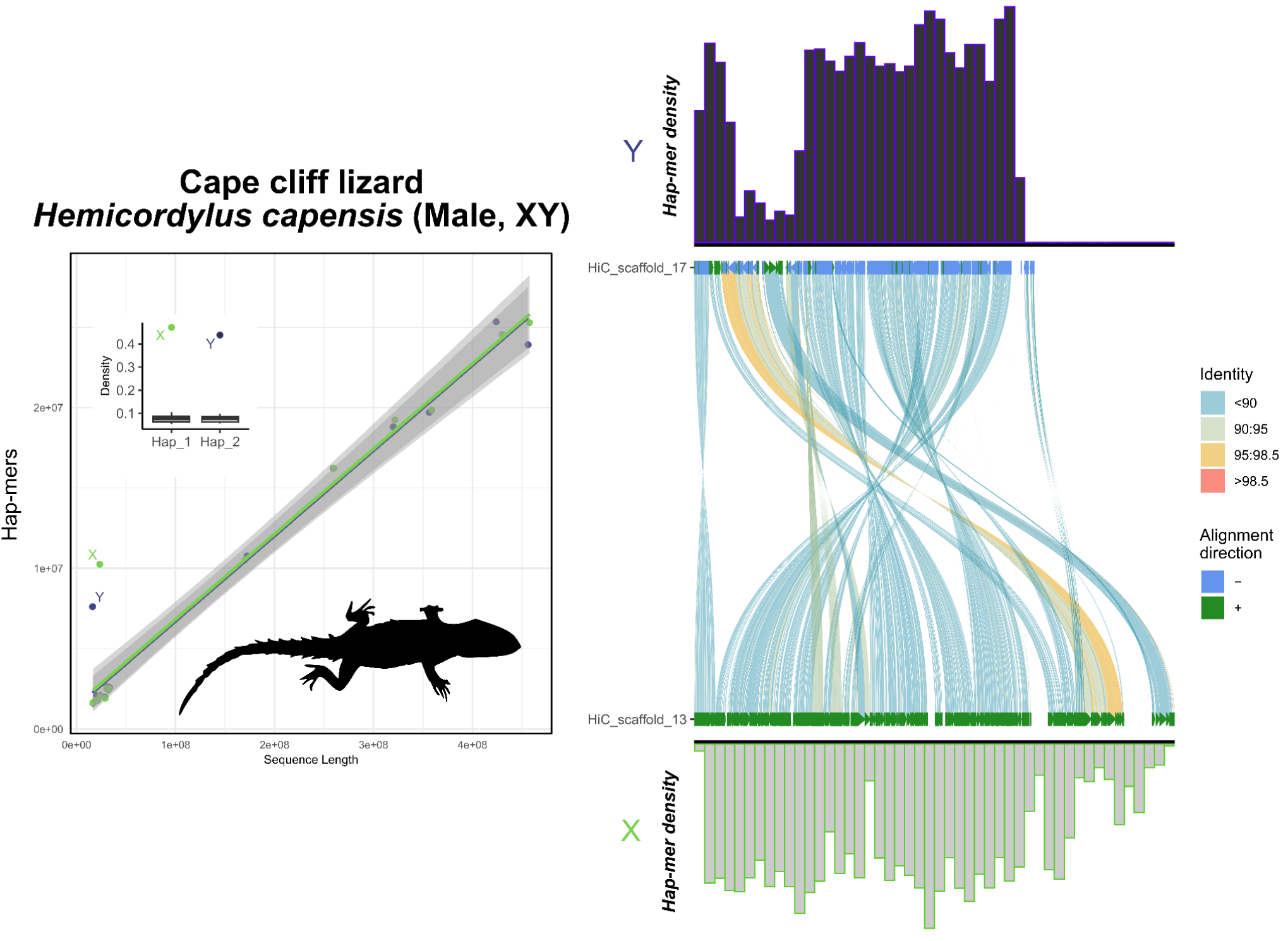
Total evidence plot for the Cape cliff lizard, *Hemicordylus capensis*, under the SCINKD framework. The left panel consists of SCINKD “results” output ∼ sequence length (Li et al., 2009) visualized with ggplot2 (Wickham, 2016). The right panel corresponds to a minimap2 alignment (Li, 2018) of putative sex chromosome haplotypes visualized using SVbyEye (Porubsky et al., 2024) overlaid with hap-mer densities in 1Mb windows across the chromosome. A single pair of haplotypes deviates from the autosomal background. The signal comes from nearly the entirety of LG13, a homologous pair that also aligns with relatively low sequence identity across its length. There is no evidence that the PAR region is present in this assembly. The convergence of evidence on a single region corroborates previous work showing this is the XY linkage group in this scincomorph lizard species (Leitão et al., 2023).

**Figure 4:**
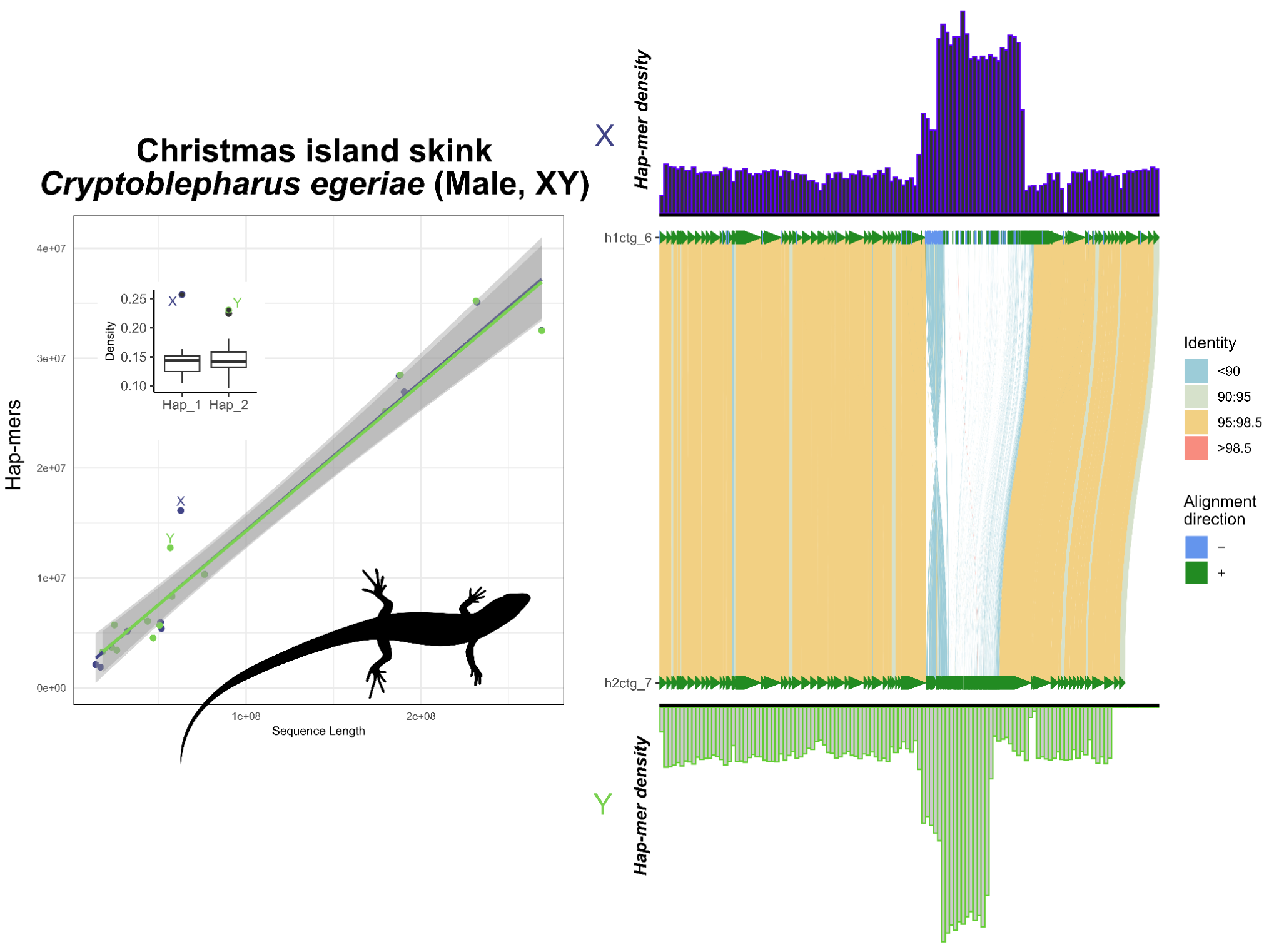
Total evidence plot for the Christmas Island skink, *Cryptoblepharus egeriae*, under the SCINKD framework. The left panel consists of SCINKD “results” output ∼ sequence length (Li et al., 2009) visualized with ggplot2 (Wickham, 2016). The right panel corresponds to a minimap2 alignment (Li, 2018) of putative sex chromosome haplotypes visualized using SVbyEye (Porubsky et al., 2024) overlaid with hap-mer densities in 1Mb windows across the chromosome. A single pair of haplotypes deviate from the autosomal background. The signal itself originates from a single region at the center of two homologous contigs, a region that aligns poorly between haplotypes and possesses an excess of unique kmers. The convergence of evidence on a single region corroborates previous work in skinks and supports this region as the XY linkage group in this scincomorph lizard species (Kostmann et al., 2021) and was validated via alignment to chrX of the Three-lined skink, *Acritoscincus duperreyi*, assembly (GCA_041722995.2).

### Conceptual framework overview

Deeper interpretation of exemplar taxa in Figure 2 among others—also supported by myriad embargoed datasets—led to a refined conceptual framework described in Figure 1. Briefly, in regions of restricted recombination, degeneration of the sex-limited chromosome is expected due to lack of germline DNA repair and evolutionarily driven by selection and drift (Bergero & Charlesworth, 2009). The absence of robust repair mechanisms causes genetic divergence between the gametologous chromosomes (between X and Y or Z and W), and this degeneration process can happen over relatively short evolutionary timeframes. Although difficult to validate in the absence of additional data (i.e. genomic information from multiple females and males), these differences often occur in excess relative to the autosomal background (Supplemental Figures 5 and 6). In heteromorphic systems, these differences are extreme (e.g. Figure 2A,C,E; Supplemental Figure 5), and can typically also be supported by read-depth related measurements. However, most vertebrate species possess homomorphic sex chromosomes (Az et al., 2009; Hillis & Green, 1990; Schultheis et al., 2009), but these systems still possess differences in excess relative to an autosomal background (Supplemental Figure 6). Further, in certain taxa with ancient sex chromosome systems (i.e. hundreds of millions of years of reduced effective population size), the sex chromosomes in a homogametic individual (XX or ZZ) can also be identified by a diagnosed by a dearth of hap-mers (Figure 2B,D) (Wilson Sayres, 2018). Under certain conditions, there may be no excess or dearth of hap-mers present (or a correlation between chromosome length and hap-mer quantity fails to appear) and in these cases SCINKD would not be effective. These conceptual points are presented in a graphical framework in Figure 1.

### Divergent and previously unannotated test cases

When extrapolating our SCINKD conceptual framework across new data, we targeted a broad sample of numerous independent sex chromosome systems with varying levels of empirical support (Figures 3-10).

1. In the crested gecko (*Correlophus ciliatus*), our results support the previously identified ZW system using hap-mer densities, a system supported by RADseq data of multiple male and female samples (Figure 5) (Gamble et al., 2015; Keating, 2022).
2. In the lesser electric ray (*Narcine bancroftii*), we expanded further across the phylogeny to observe strong support for the presumed ancestral XY linkage group on LG12. This species was assumed to be XX/XY but due to ambiguity from other methods, no sex chromosomes have been previously annotated (Figure 6) (Lee et al., 2025).
3. In the San Diegan legless lizard (*Anniella stebbinsi*), we see strong support for patterns consistent with a ZW system on LG7 (Figure 7). This is consistent with other data suggesting that anguimorphs possessed an ancestral ZZ/ZW sex chromosome system with multiple transitions to novel ZW linkage groups (Pinto et al., 2024; Rovatsos et al., 2019).
4. In the Christmas Island gecko (*Lepidodactylus listeri*), we see low support for patterns consistent with an XY system on LG18 (Figure 8). Relatively little data is available across this extraordinarily diverse family, Gekkonidae, to assess the likelihood or conservation of a putative XX/XY system in this gecko species. However, it’s recently been hypothesized that *Lepidodactylus* possess a ZZ/ZW system, suggesting a misassembly on LG18 is likely the cause of this signal (Gamble, 2010; Volobouev & Pasteur, 1988).

**Figure 5:**
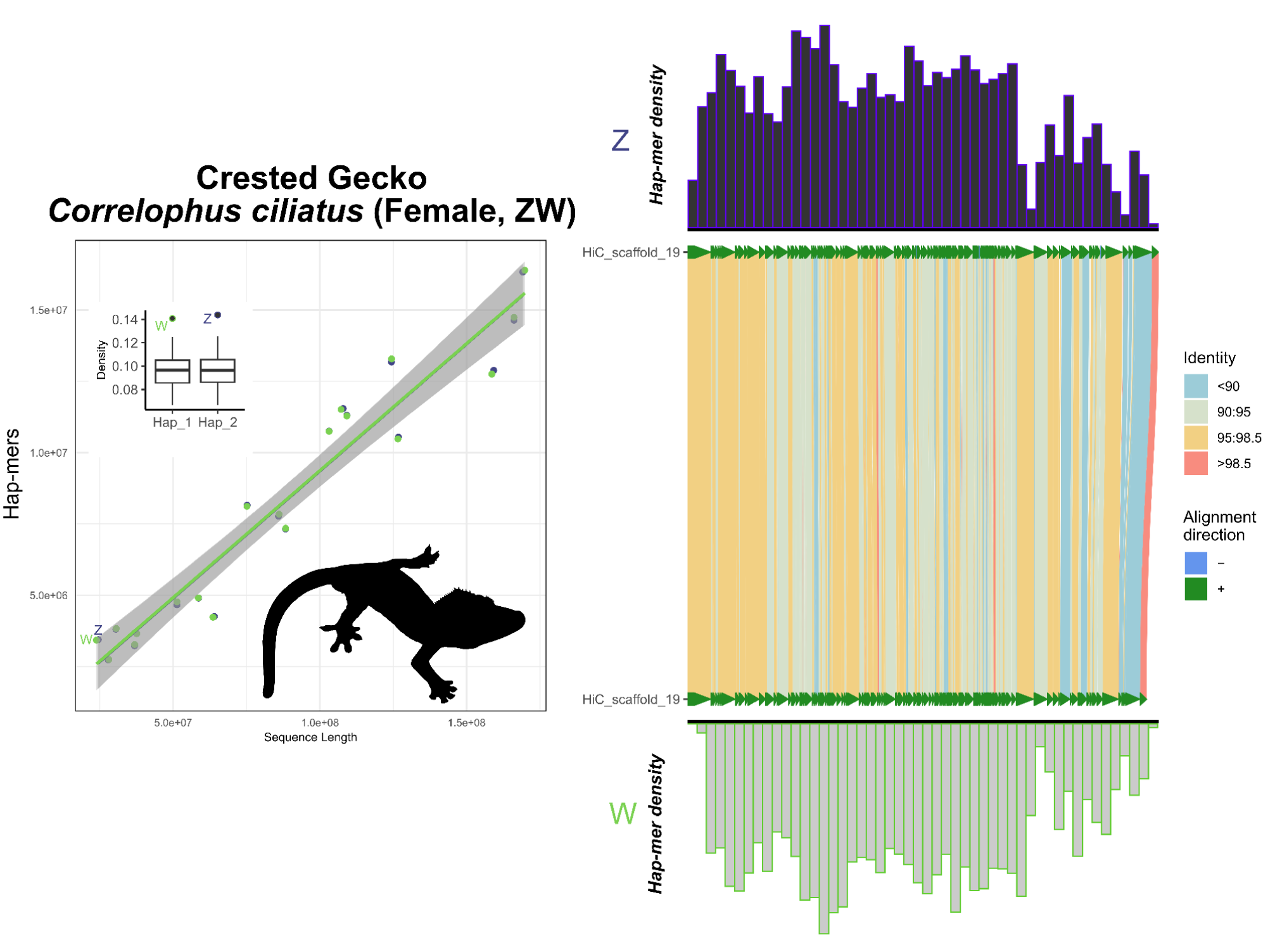
Total evidence plot for crested gecko, *Correlophus ciliatus*, under the SCINKD framework. The left panel consists of SCINKD “results” output ∼ sequence length (Li et al., 2009) visualized with ggplot2 (Wickham, 2016). The right panel corresponds to a minimap2 alignment (Li, 2018) of putative sex chromosome haplotypes visualized using SVbyEye (Porubsky et al., 2024) overlaid with hap-mer densities in 1Mb windows across the chromosome. A single pair of haplotypes deviates from the autosomal background. The signal itself originates from LG19 across most of their length, a chromosomal pair that possesses generally low sequence identity across its length with the exception of a small region at the distal tip, consistent with a pseudoautosomal region (PAR). This evidence complements and is confirmed by previous work showing that LG19 is the ZW linkage group in the crested gecko and annotated the Z and W chromosomes using sex-specific RADtags (Gamble et al., 2015; Keating, 2022).

**Figure 6:**
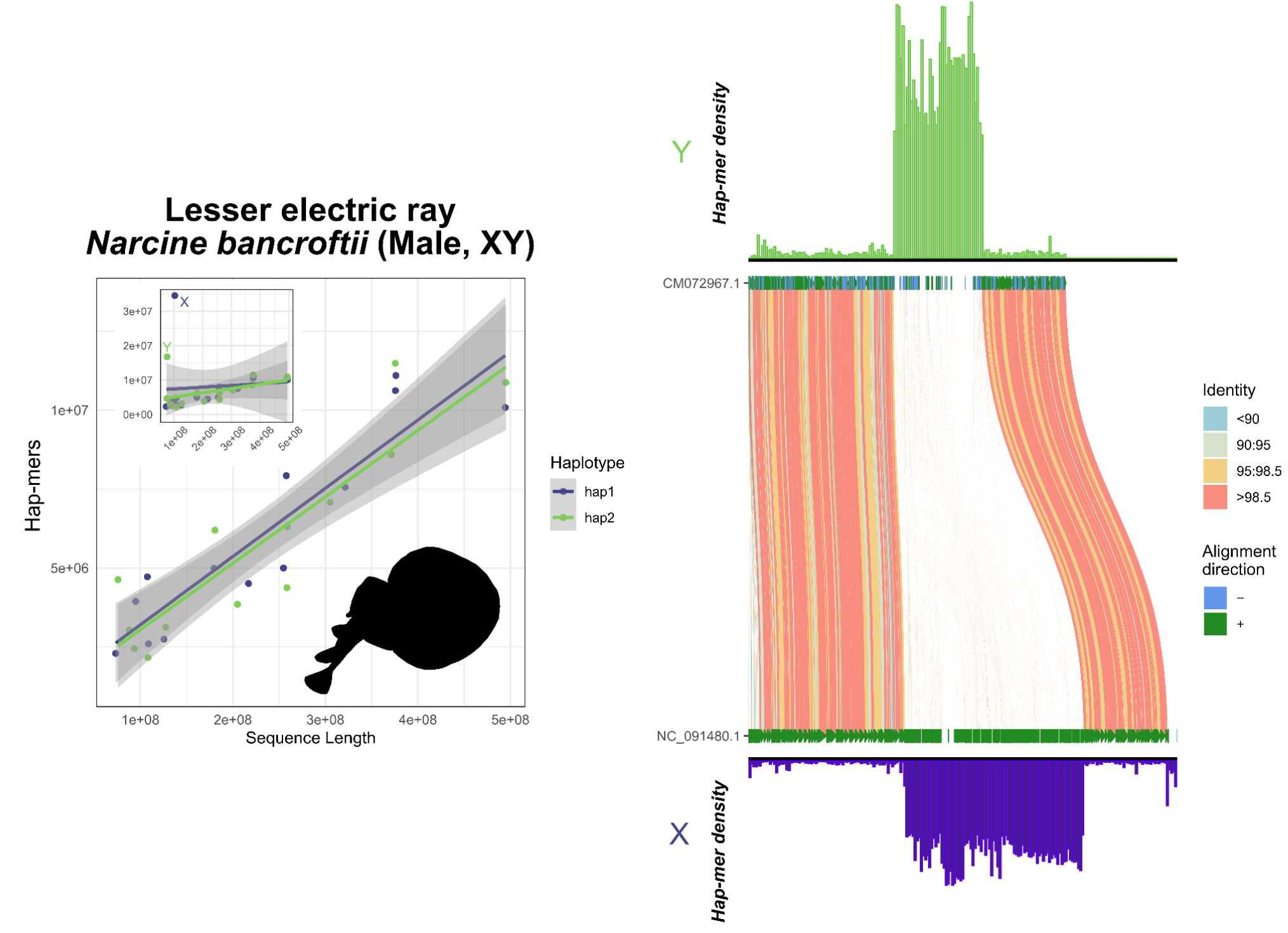
Total evidence plot for the lesser electric ray, *Narcine bancroftii*, under the SCINKD framework. The left panel consists of SCINKD “results” output ∼ sequence length (Li et al., 2009) visualized with ggplot2 (Wickham, 2016). The right panel corresponds to a minimap2 alignment (Li, 2018) of putative sex chromosome haplotypes visualized using SVbyEye (Porubsky et al., 2024) overlaid with hap-mer densities in 1Mb windows across the chromosome. A single pair of haplotypes deviates from the autosomal background. The signal itself originates from a single region at the center of LG12, a region that aligns poorly between haplotypes and possesses an excess of unique kmers. The convergence of evidence on this region corroborates previous work showing this is the conserved XY linkage group in many elasmobranch species (Lee et al., 2025). Here, the X and Y are annotated based on homology with previously annotated X and Y in related species.

**Figure 7:**
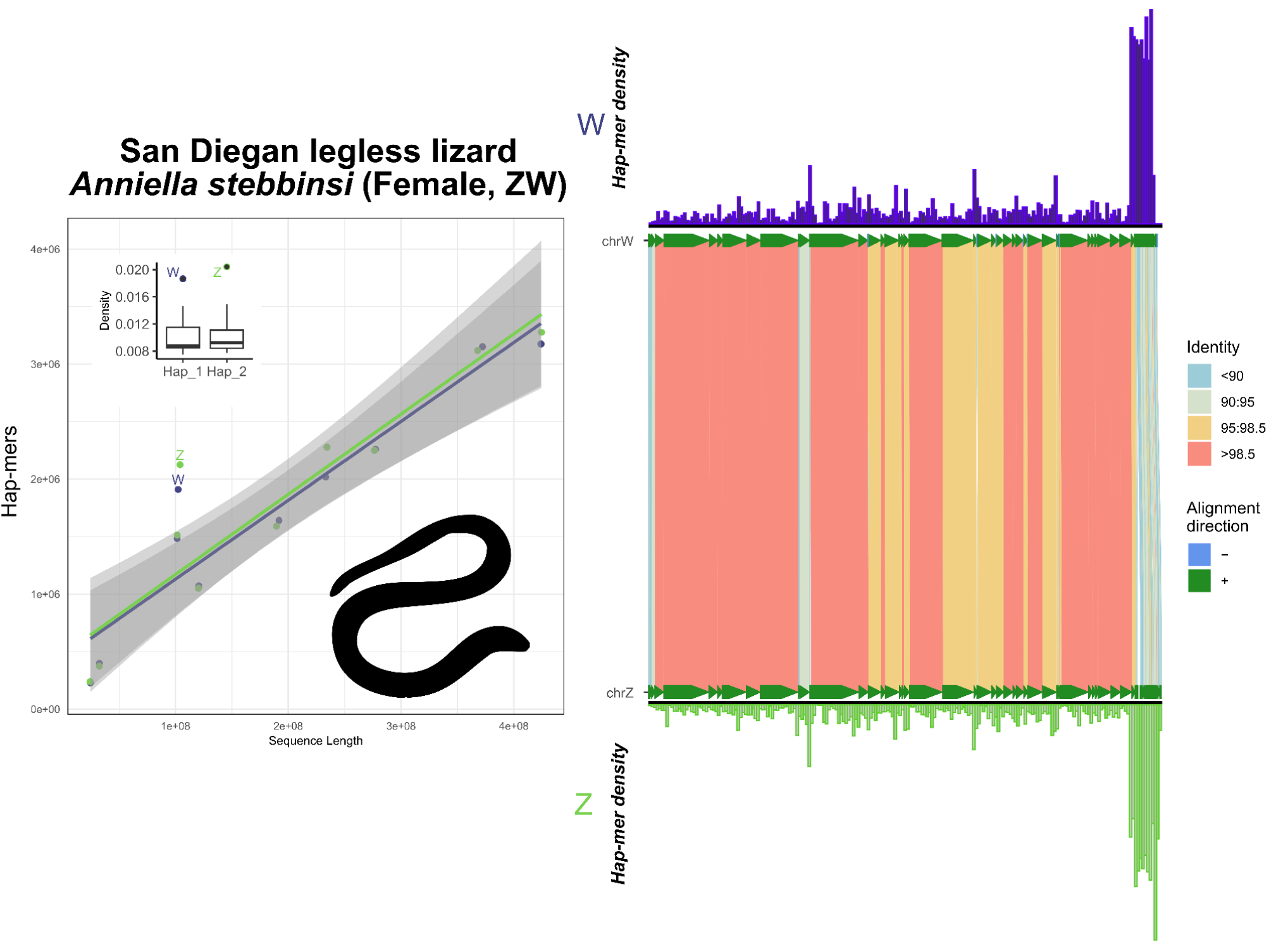
Total evidence plot for the San Diegan legless lizard, *Anniella stebbinsi*, under the SCINKD framework. The left panel consists of SCINKD “results” output ∼ sequence length (Li et al., 2009) visualized with ggplot2 (Wickham, 2016). The right panel corresponds to a minimap2 alignment (Li, 2018) of putative sex chromosome haplotypes visualized using SVbyEye (Porubsky et al., 2024) overlaid with hap-mer densities in 1Mb windows across the chromosome. A single pair of haplotypes deviates from the autosomal background. The signal itself originates from a single region at the distal end of LG7, a region that possesses reduced sequence identity relative to the rest of the pairwise alignment. The convergence of evidence on a single region provides strong support for a hypothesis of a ZZ/ZW system in this anguimorph lizard. Here, the Z and W were hypothesized due to sequence length and additionally supported by alignment to the corresponding linkage group (NC_086184.1) in the Southern alligator lizard, *Elgaria multicarinata*, assembly (GCF_023053635.1), where average sequence identity was slightly higher on the putative Z-specific region (81%) than on the W-specific region (80%), which would expected due to higher rates of sequence divergence on chrW.

**Figure 8:**
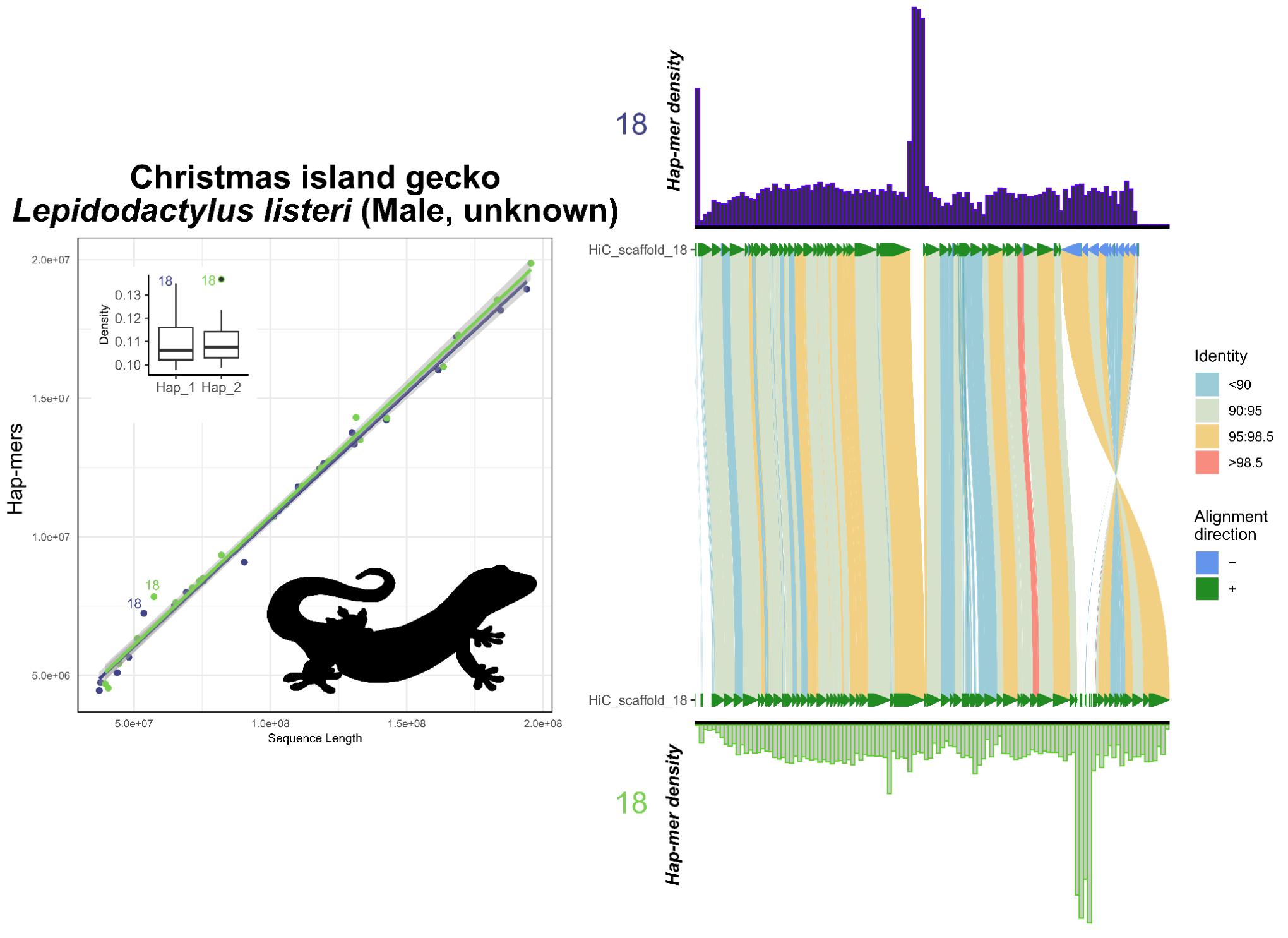
Total evidence plot for the Christmas island gecko, *Lepidodactylus listeri*, under the SCINKD framework. The left panel consists of SCINKD “results” output ∼ sequence length (Li et al., 2009) visualized with ggplot2 (Wickham, 2016). The right panel corresponds to a minimap2 alignment (Li, 2018) of putative sex chromosome haplotypes visualized using SVbyEye (Porubsky et al., 2024) overlaid with hap-mer densities in 1Mb windows across the chromosome. A single pair of haplotypes deviates from the autosomal background. The signal itself originates from two poorly aligned regions between the haplotype pairs. The convergence upon a linkage group of interest, but not focused on a specific region, provides relatively low support for a hypothesis that LG18 is the XY linkage group in this gecko species, but could warrant further investigation using gene annotations and data from additional individuals. Low confidence of support for this putative XY system preempts any attempt to assign haplotypes as putative X and Y.

Neither latter species, *A. stebbinsi* and *L. listeri*, has publicly available resequencing data from males and females to further validate these hypotheses. However, in cases like these where additional data is unavailable or prohibitive, our results suggest SCINKD is more robust than a read depth-related approach in supporting candidate sex-linked regions in recently evolved sex chromosome systems (Pinto et al., 2022).

Next, to examine SCINKD robustness using a more diverse taxon, we tested SCINKD with an attempt to independently confirm the sex chromosome linkage group in a New Caledonia shrub species, *Amborella trichopoda*, with known ZZ/ZW sex chromosomes (Carey et al., 2024). In this shrub, we observe strong deviations from our vertebrate-focused model of correlations between hap-mers and chromosome length (Figure 9). However, the ZW chromosomal pair still possess the most hap-mers. The lack of correlation between hap-mers and chromosome length can likely be attributed to the distinct mechanisms of recombination present in plants (Gaut et al., 2007), which is thought to drive much of the previously described correlations of chromosome length in vertebrates (Pinto, Gamble, Smith, & Wilson, 2023). Regardless, since hap-mer densities appear to be disproportionate on the sex chromosomes in *Amborella*, there may be potential for developing a more broad SCINKD conceptual model that accounts for alternative biology present in non-vertebrate lineages (Figure 9).

**Figure 9:**
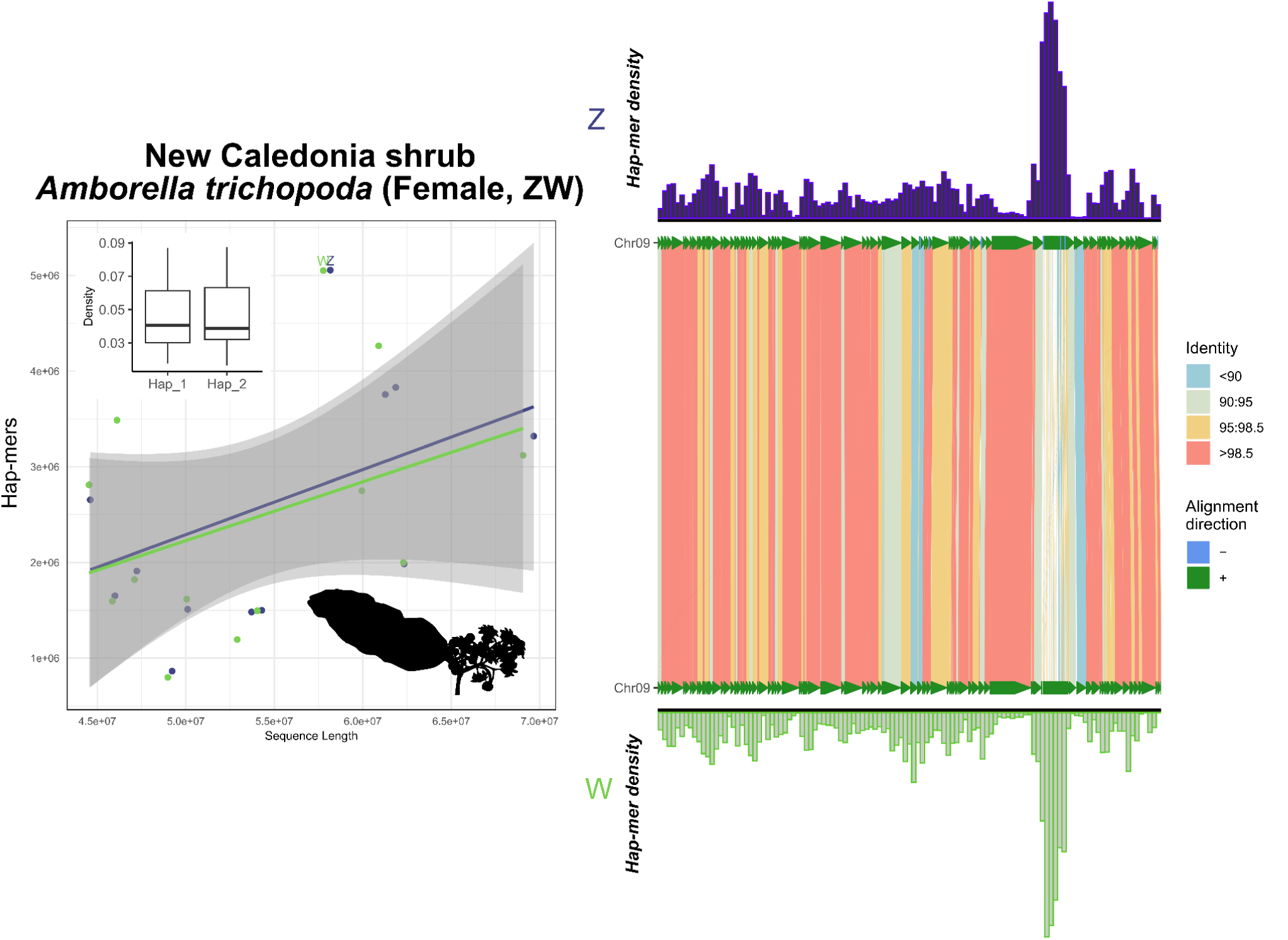
Total evidence plot for the New Caledonia shrub, *Amborella trichopoda*, under the SCINKD framework. The left panel consists of SCINKD “results” output ∼ sequence length (Li et al., 2009) visualized with ggplot2 (Wickham, 2016). The right panel corresponds to a minimap2 alignment (Li, 2018) of putative sex chromosome haplotypes visualized using SVbyEye (Porubsky et al., 2024) overlaid with hap-mer densities in 1Mb windows across the chromosome. A single pair of haplotypes deviates from the autosomal background. The signal itself originates from a single region at the distal end of LG7, a region that possesses reduced sequence identity relative to the rest of the pairwise alignment. The haplotype density and pairwise alignment in the previously identified sex-limited region is concordant with previous work showing these are the Z and W chromosomes in this species of shrub (Carey et al., 2024). However, no robust statistical support is observed under the SCINKD framework, suggesting plants do not conform to our vertebrate-centric linear predictions of haplotype divergence.

**Figure 10:**
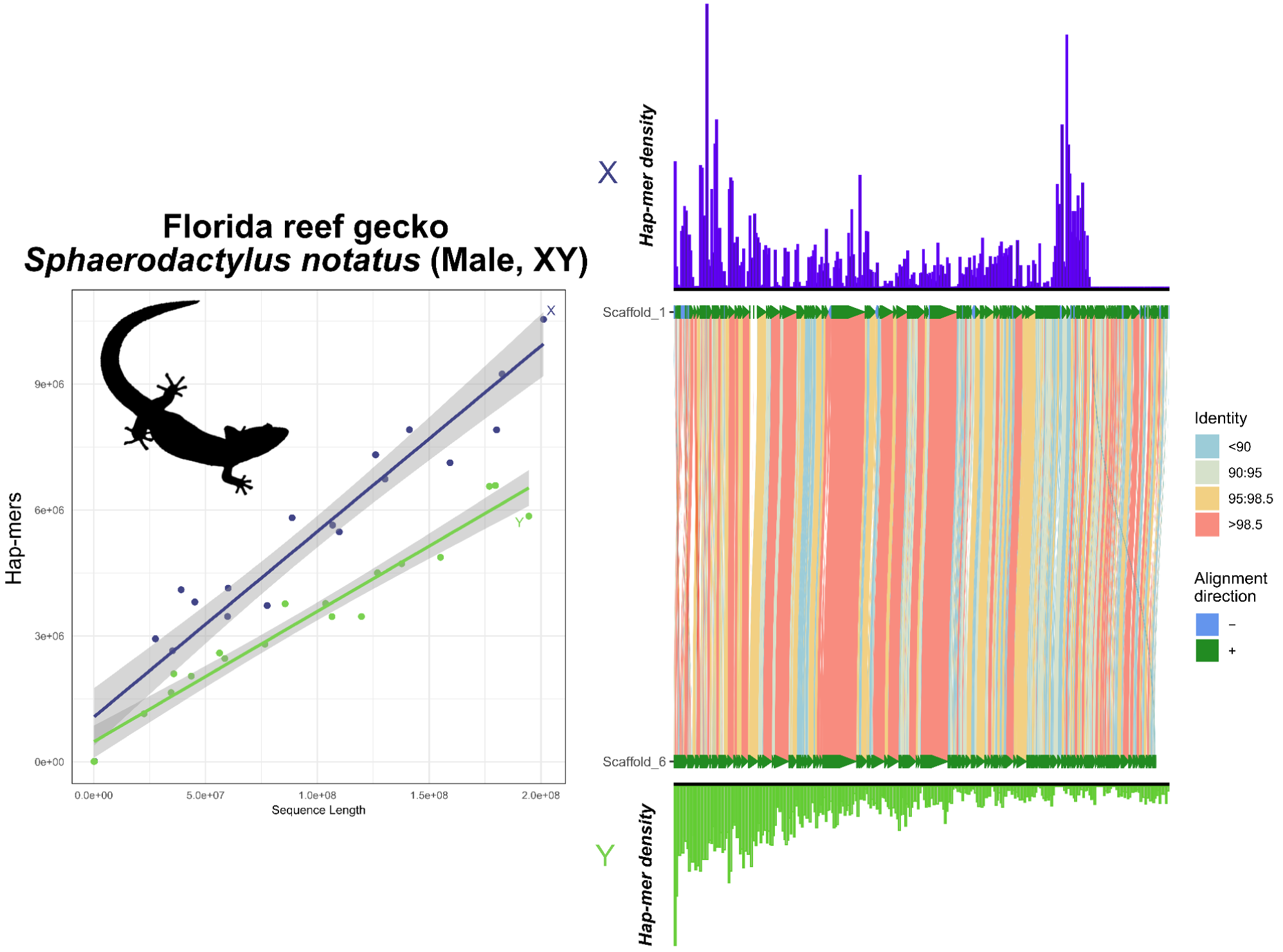
Total evidence plot for the Florida reef gecko, *Sphaerodactylus notatus*, under the SCINKD framework. The left panel consists of SCINKD “results” output ∼ sequence length (Li et al., 2009) visualized with ggplot2 (Wickham, 2016). The right panel corresponds to a minimap2 alignment (Li, 2018) of putative sex chromosome haplotypes visualized using SVbyEye (Porubsky et al., 2024) overlaid with hap-mer densities in 1Mb windows across the chromosome. We see no explicit evidence of the XX/XY system in this species under the SCINKD framework. A lack of convergence upon a common slope suggests phasing issues with the genome assembly. The sex chromosome linkage group itself, identified by previous work (Pinto et al., 2022), also shows no evidence of sex chromosome patterns in the sex-limited region suggesting an excess of noise, likely caused by a combination of low sequencing coverage (∼20x) and a dearth of heterozygosity. However, we were able to annotate the X and Y chromosomes using previously published sex-specific RADtags (Pinto et al., 2022).

Lastly, we developed an opportunistic test case for a difficult assembly with a known XX/XY sex chromosome system in the Neotropical leaf-litter gecko, *Sphaerodactylus notatus* (Pinto et al., 2022). In this experiment, the *S. notatus* sample contained technical artifacts from a combination of (1) low HiFi sequencing yield (∼20x coverage), (2) shorter-than-optimal read lengths (mean read length=∼13.5kb), and (3) a dearth of heterozygosity (0.35%) perhaps due to its hypothesized low effective population size (Clements et al., 2021)—which is even below the rate previously documented to cause phasing issues in other taxa propagated in the pet trade, e.g. 0.56% in *Eublepharis macularius* (Pinto, Gamble, Smith, Keating, et al., 2023).

Under these conditions, we see our first instance of a vertebrate deviating from expectations (Figure 10). Here, chr1 is the known XY linkage group (Pinto et al., 2022). However, low coverage and technical artifacts prevent the haplotypes from conforming to the expected correlation between sequence length to hap-mer abundance and preempts SCINKD from isolating them. Indeed, because the SCINKD framework is a biological prediction, the diploid reference genome data representing that biology may possess inherent biases that can inhibit genome assembly and/or phasing on the sex chromosomes (Antipov et al., 2024; Cheng et al., 2022; Pinto, Gamble, Smith, Keating, et al., 2023; Rhie et al., 2020). Thus, because there is a direct effect of sequencing depth on per-base quality and completeness of any given genome assembly, insufficient coverage may be a recurrent issue in many species.

### What does SCINKD support mean?

SCINKD was specifically designed to identify homomorphic sex chromosome systems. Although by extension it works well in heteromorphic systems there are other robust methods developed to identify heteromorphic sex chromosomes, including cytogenetics and read depth analysis (https://github.com/makovalab-psu/HalfDeep). SCINKD provides a multidimensional breakdown of haplotype uniqueness that helps inform the likelihood of sex-limited regions in the genome. At a global scale (dot plots and box plots), we develop a robust framework to investigate statistical outliers (e.g. Figure 7). After a candidate(s) are identified, one can assess hap-mer density on a finer scale using mirror plots (e.g. Supplemental Figures 3,4,5). Lastly, corroborating these layers with 1:1 chromosome alignments for candidate linkage groups (e.g. using minimap2 and SVbyEye) provides a link between chromosome coordinates and haplotype-specific anomalies (Figure 7). Indeed, in *A. stebbinsi*, a homomorphic system we view as providing strong evidence under this framework, each of the aforementioned analyses converge on a single set of haplotypes where hap-mer density increases exponentially and sequence identity between haplotypes drops sharply (Figure 7). These factors taken together provide evidence to hypothesize a ZZ/ZW system in *A. stebbinsi*.

### Caveats and limitations

We hypothesize that phasing issues may be the primary cause for excessive noise observed under the proposed framework. We show here that sequencing depth itself, above a reasonable threshold (>20x), does not greatly affect sex chromosome signals in the high-quality sequencing data for the putatively homomorphic ZW system in *Anniella stebbinsi* (Supplemental Figure 2). However, misassemblies as small as 1Mb may begin to obscure sex chromosome signals (Supplemental Figure 1). Thus, as long as a sequencing depth and heterozygosity are great enough to accurately phase the genome, SCINKD may be an exceedingly useful tool for sex chromosome identification. We show that per-haplotype sequencing depth needs to exceed the standard of 10x coverage using PacBio HiFi reads to unobscure sex chromosome signals in our *Anniella stebbinsi* example (Supplemental Figure 2). However, greater sequencing depth reduces errors and can help prevent misphasing and collapses, which can obscure otherwise clear signals from the sex chromosomes (Supplemental Figure 1). It remains untested how equivalent ONT data performs in this context. A more robust sensitivity analysis in the future is warranted when a broad taxonomic sampling of high-coverage ONT data is publicly available. Preliminary data shared with us across diverse taxa suggest that 30x ONT coverage may not be sufficient to prevent the aforementioned technical artifacts that interfere with the underlying assumptions of the SCINKD framework (A. Barley & L. Gray *pers. comm.*), although 100x coverage is more than sufficient to identify the heteromorphic sex chromosome system in the Gila monster (*unpublished data*). In sum, to help address technical artifacts that interfere with haplotype resolution across the genome, but especially the sex chromosomes where the sex-limited region is assembled largely independently, we believe recommending coverage at 20-30x per-haplotype (40-60x total coverage) is justifiable to assist in identification of sex chromosomes in many diverse taxa. This recommendation coincides with recent increases in sequencing outputs and deviates from the current recommended standards of 30x total coverage for a typical vertebrate assembly (Pinto, Gamble, Smith, & Wilson, 2023).

The data do not yet currently exist to characterize every possible outcome across divergent taxa, technical capacity, or sex chromosome system. Observations presented here also suggest that the Y/W typically falls lower on both axes of the dot plot, irrespective of chromosome length. This is presumably due to (1) the sex-limited chromosome (W or Y) being typically shorter than its gametologous counterpart, and (2) accumulation of repetitive DNA and loss of gene content, both of which can reduce the numbers of unique hap-mers on the Y/W relative to their X/Z homolog, and relative to their own length. However, as is often the case in biology, this is not always true and should not be used as the only metric for sex chromosome assignment. Another notable observation here is that no two sex chromosome alignments share any notable feature that would independently indicate assignment as a sex chromosome (Figure 2-10), which is why using a pairwise alignment alone to identify candidate loci across diverse systems is not sufficient. Thus, given the theoretical framework presented here (Figure 1) and the ability to view multivariate outputs from the SCINKD workflow (i.e. dotplots, density boxplots, mirror plots), combined with pairwise alignments, we provide researchers the ability to make strong hypotheses regarding sex chromosome assignments across diverse taxa from a single, diploid genome. We emphasize the importance of an empirical approach (an understanding of the biology of the study system as well as sex chromosome evolution) when applying this framework.

### How should these sex chromosomes be annotated?

It’s worth mentioning a few important observations of the data presented here. No individual analysis (e.g. HiC data, read depth, haplotype alignment, or SCINKD results) can be taken alone as conclusive evidence that a pair of haplotypes make up a novel sex chromosome system. Indeed, when applying methods such as SCINKD at scale, we will continue to provide strong evidence for novel putative sex chromosomes where no other data is available for validation, such as in the San Diegan legless lizard (Figure 7). We suggest a 4-category system for sex chromosome annotation for any particular genome, and cluster taxa from this study into them as examples. We describe two broad scenarios, differentiated by levels of available evidence, but always recommend cross-referencing using data from additional individuals when available.

1. In species with strong evidence of sex chromosomes, the sex chromosomes should be annotated, (A) the “gold standard”—those validated by external data types (such as resequencing data, RADseq, or a high-quality assembly of both sexes, etc)—e.g. mammals/birds/lacertids (Figure 2), cape cliff lizard (Figure 3) (Leitão et al., 2023), crested gecko (Figure 5) (Gamble et al., 2015), New Caledonia shrub (Figure 9) (Carey et al., 2024), and Florida reef gecko (Figure 10) (Pinto et al., 2022), or (B) corresponding evidence from SCINKD and another source(s) (such as phylogenetic history and/or read depth from the assembly input data)—e.g. Christmas island skink (Figure 4), lesser electric ray (Figure 6), and San Diegan legless lizard (Figure 7).
2. In species with moderate-to-low evidence of sex chromosome linkage information, the sex chromosomes should *not* be annotated and await additional data (such as resequencing data, a high-quality assembly of the opposite sex, RADseq, etc), such as (C) those assemblies with a statistically predicted candidate linkage group with no candidate sex determining region or phylogenetic information—e.g. the Christmas island gecko (Figure 8), or (D) those with no clear evidence of a sex chromosome linkage group in the assembly—e.g. the ZZ Nicobar pigeon (Figure 2D), ZZ Cretan wall lizard (Figure 2F), or the TSD leopard gecko (Supplemental Figure 4).

Here, we present myriad examples from typical vertebrate (and a plant) sex chromosome systems with varying levels of success. However, sex determination itself is a complicated process and many species do not even possess sex chromosomes. For example, given that hymenopterans possess complementary sex determination (CSD), we can confidently state that well-over 20% of total extant animal species do not possess sex chromosomes (Heimpel & de Boer, 2008; Whiting, 1943). Outside of Metazoa, approximately 5-10% of angiosperms use genetic sex determination (Charlesworth & Harkess, 2024), but to our knowledge *A. trichopoda* is the only plant species to date with assembled sex chromosomes in a haplotype-resolved assembly. Beyond that, other aspects of life history vary wildly across taxa, from TSD to polygenic sex determination to species utilizing a sex-limited mitochondrial region as a locus of large effect (Kocher et al., 2024; Smith et al., 2023). In any of these instances, it is possible that SCINKD may assist in genome curation but would be improperly utilized to hypothesize the existence of sex chromosomes in these types of diverse taxa. Thus, since the null predictions used in SCINKD are biological with technical considerations applied to interpretation, it is important to understand the biology of each system prior to generating any hypotheses using SCINKD.

## Conclusion

The utility of SCINKD in the age of biodiversity genomics is apparent and has already been deployed to assist with sex chromosome curation across the most recent genome assemblies from the Vertebrate Genomes Project (Rhie et al., 2021). However, there also remain vertebrate lineages that remain underrepresented by the VGP, such as squamates, the most speciose discrete clade of tetrapods (Pinto, Gamble, Smith, & Wilson, 2023; Rhie et al., 2021). As these lesser-known taxa across the tree of life begin acquiring reference genomes, there will continue to be a lack of population-level genomic data traditionally used to develop sex chromosome annotations. We view the SCINKD framework as a significant step forward towards addressing the ongoing challenges in sex chromosome assembly and annotation at a scale to meet the rapidly moving field of genomics.

## Acknowledgements

The authors would like to acknowledge M. Kirkpatrick and J. Havird for additional funding and laboratory space for the *S. notatus* HiC data and comments on the manuscript; B. Pickett for assistance in navigating meryl syntax between SCINKD versions 1 and 2; S. Plaisier for discussion on the nascent concept of SCINKD; and L. Gray and A. Barley for sharing unpublished ONT genomes and data for additional sensitivity testing. For novel data presented here, animal husbandry and tissue sampling were conducted under Marquette University IACUC protocol AR-279. The authors acknowledge Research Computing at Arizona State University for providing high performance computing resources that have contributed to the research results reported within this paper.

## Code and data availability

Both SCINKD code and user tutorial are available on GitHub (https://github.com/DrPintoThe2nd/SCINKD). Raw data generated in this study is available on SRA (SAMN48008689-90) and genome reassemblies on Figshare (https://doi.org/10.6084/m9.figshare.29344685) and, where applicable, also to GenBank.

## Author contributions

BJP*^‡^ conceptualized the project, acquired funding, developed the tool, conducted genome sequencing and assembly experiments, and drafted the manuscript; SG^‡^ ran vertebrate-wide analyses and generated data figures; SEK generated RADseq library; CHS acquired funding and assisted with HiC experiment; TG provided materials and laboratory space; SVN acquired funding, feedback, and the *Sphaerodactylus* sample; MAW* acquired funding, supervised the project, and provided regular feedback.

## Supplemental materials

**Supplemental Table 1:**
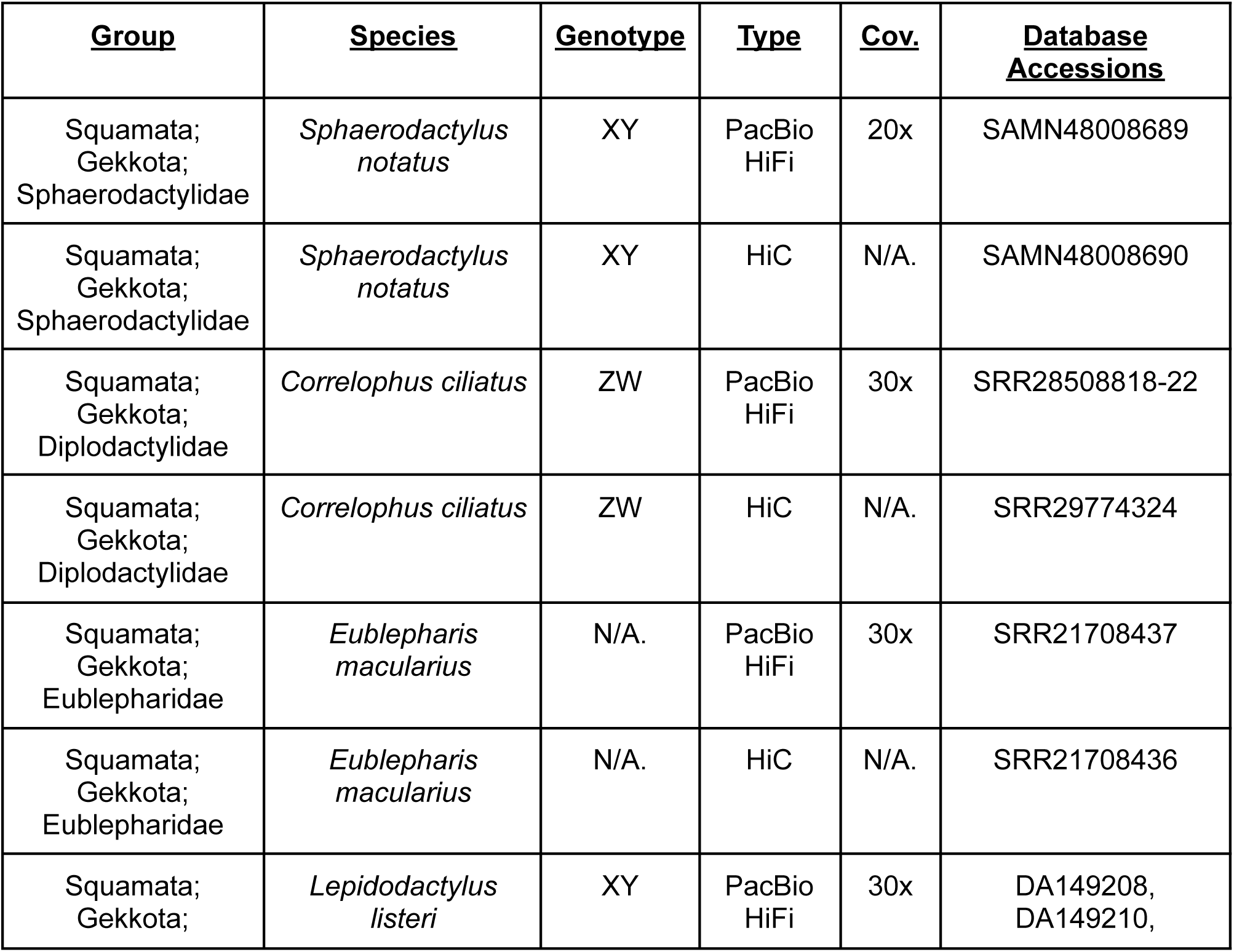
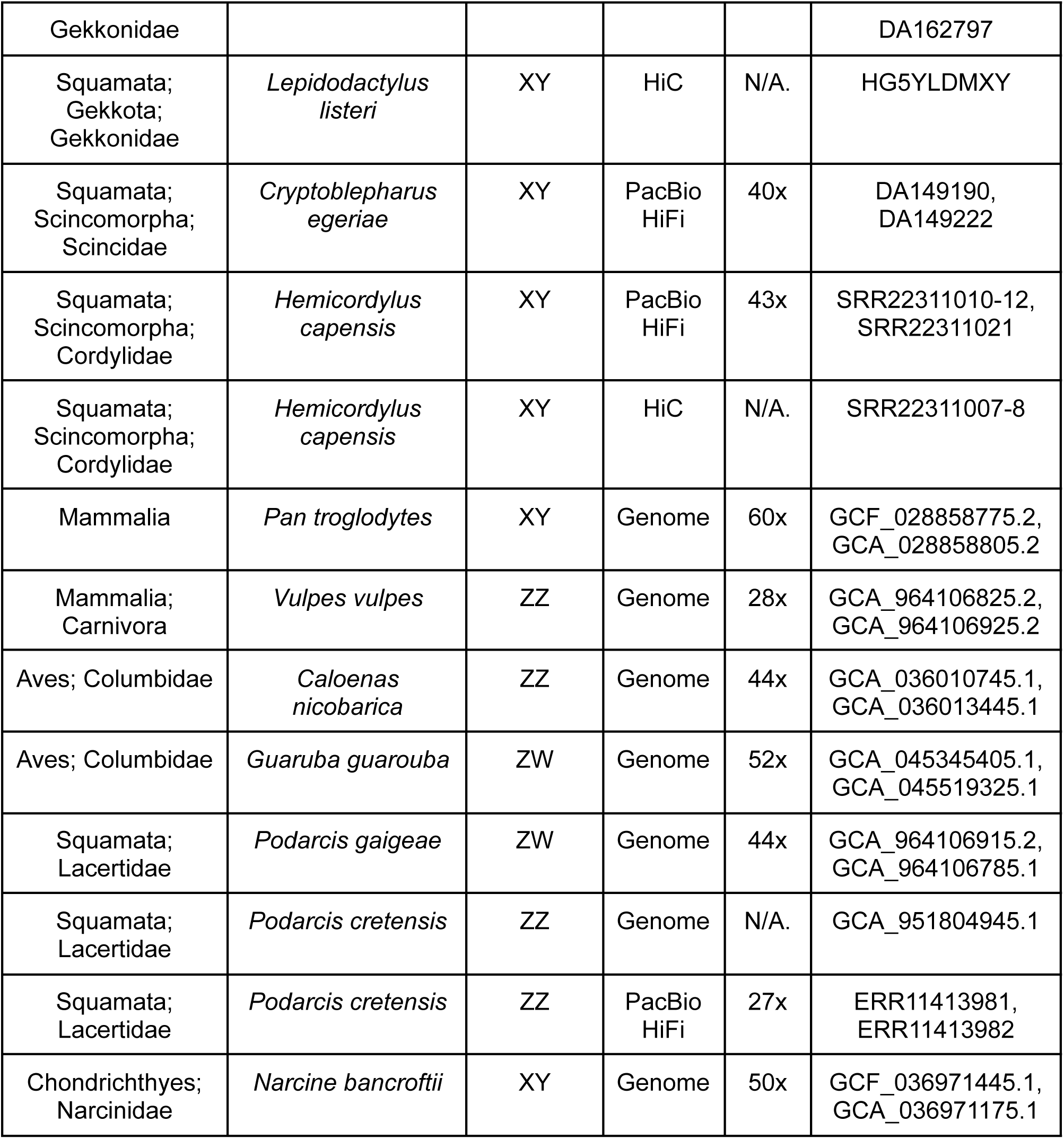
Sample list and accession numbers.

**Supplemental Figure 1:**
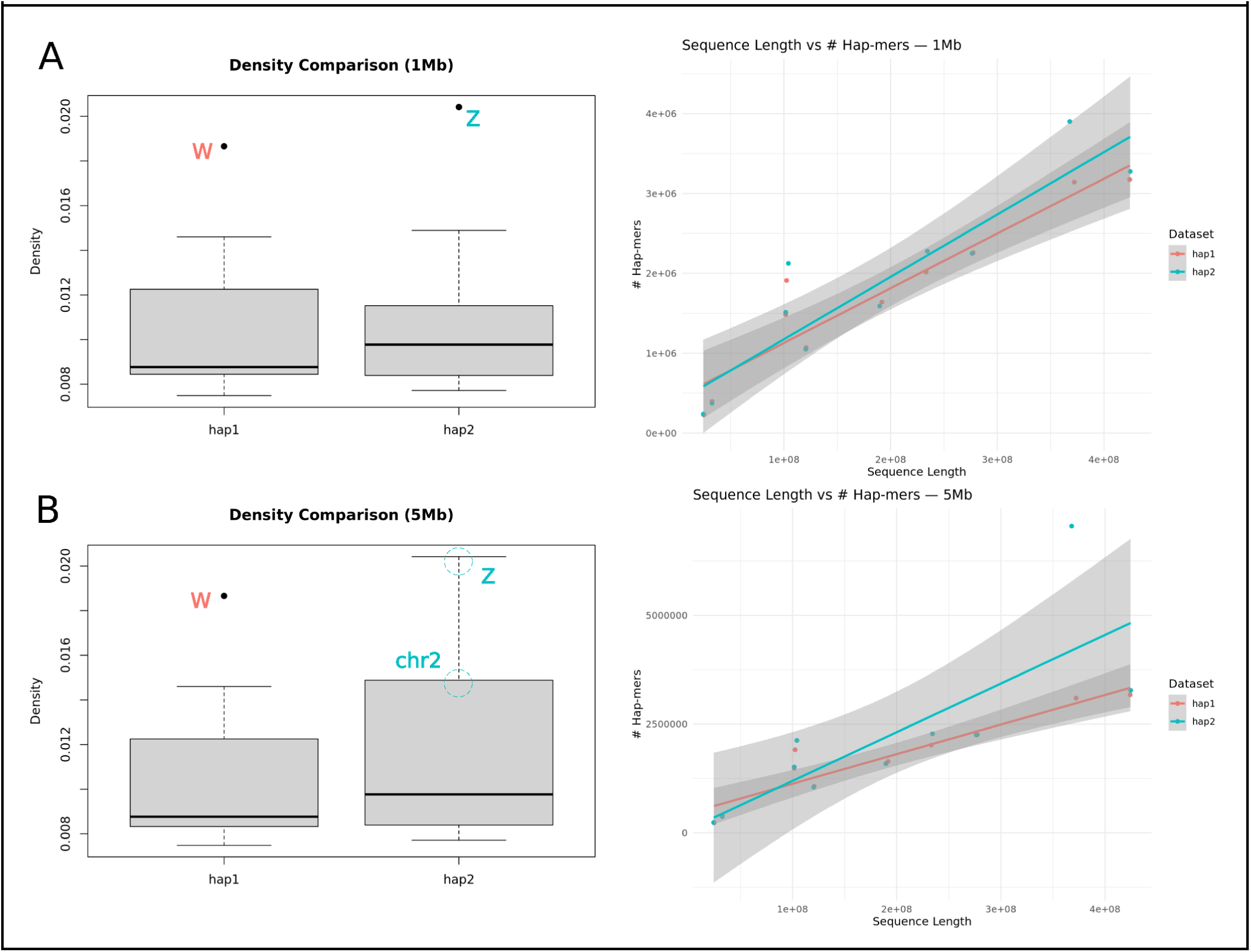
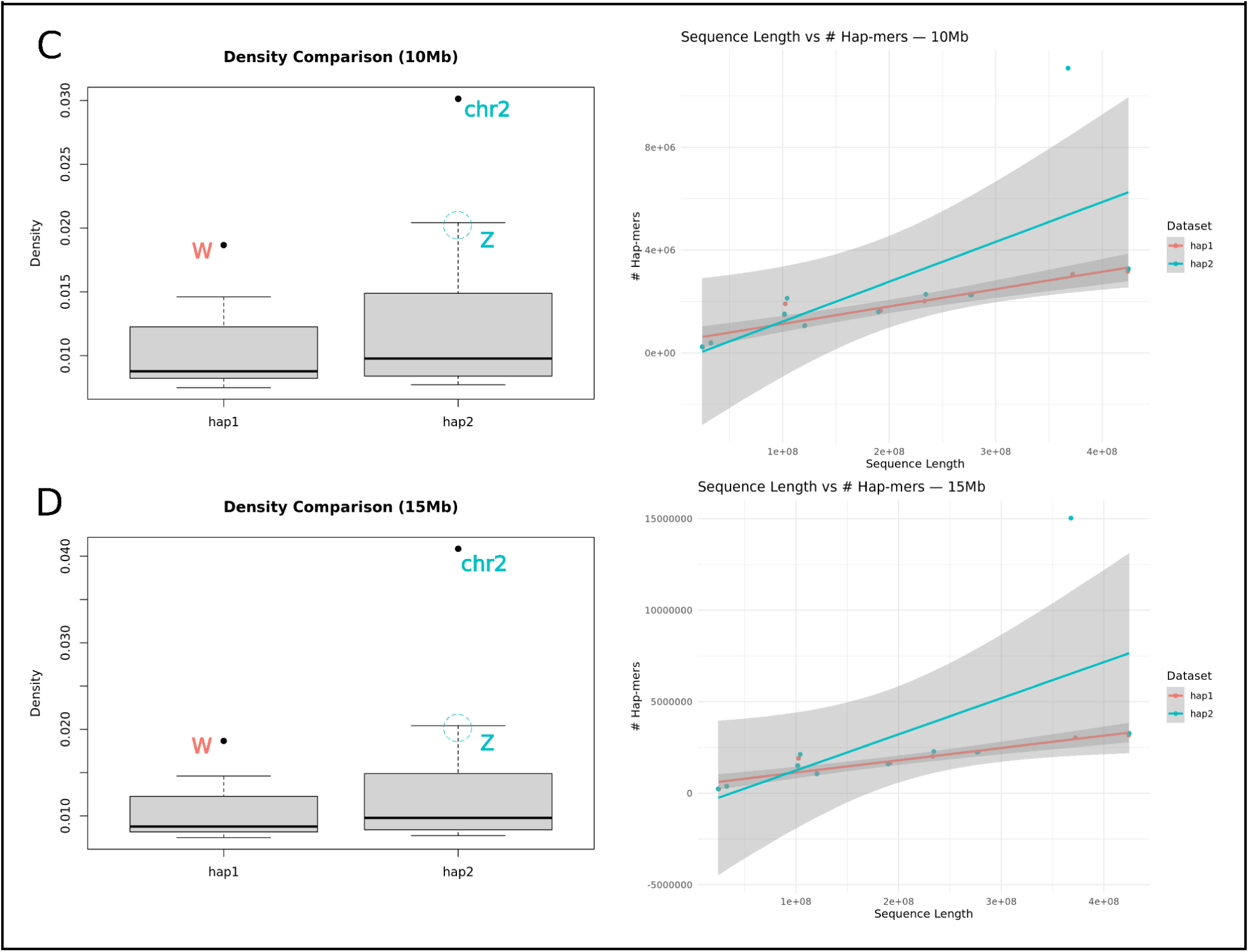
Using the updated reference assembly for Anniella stebbinsi detailed in the main text. We systematically hard-masked a region on chromosome 2 in haplotype 1 of varying sizes to mimic a collapsed homozygous region in the assembly. (A) a 1Mb region increases noise, but retains the original sex chromosome signal, (B) a 5Mb region significantly increases noise, but qualitative assessment may still anticipate the original sex chromosome signal, (C) by 10Mb the original signal is nearly completely lost, and (D) further exacerbates the effects observed in panel C.

**Supplemental Figure 2:**
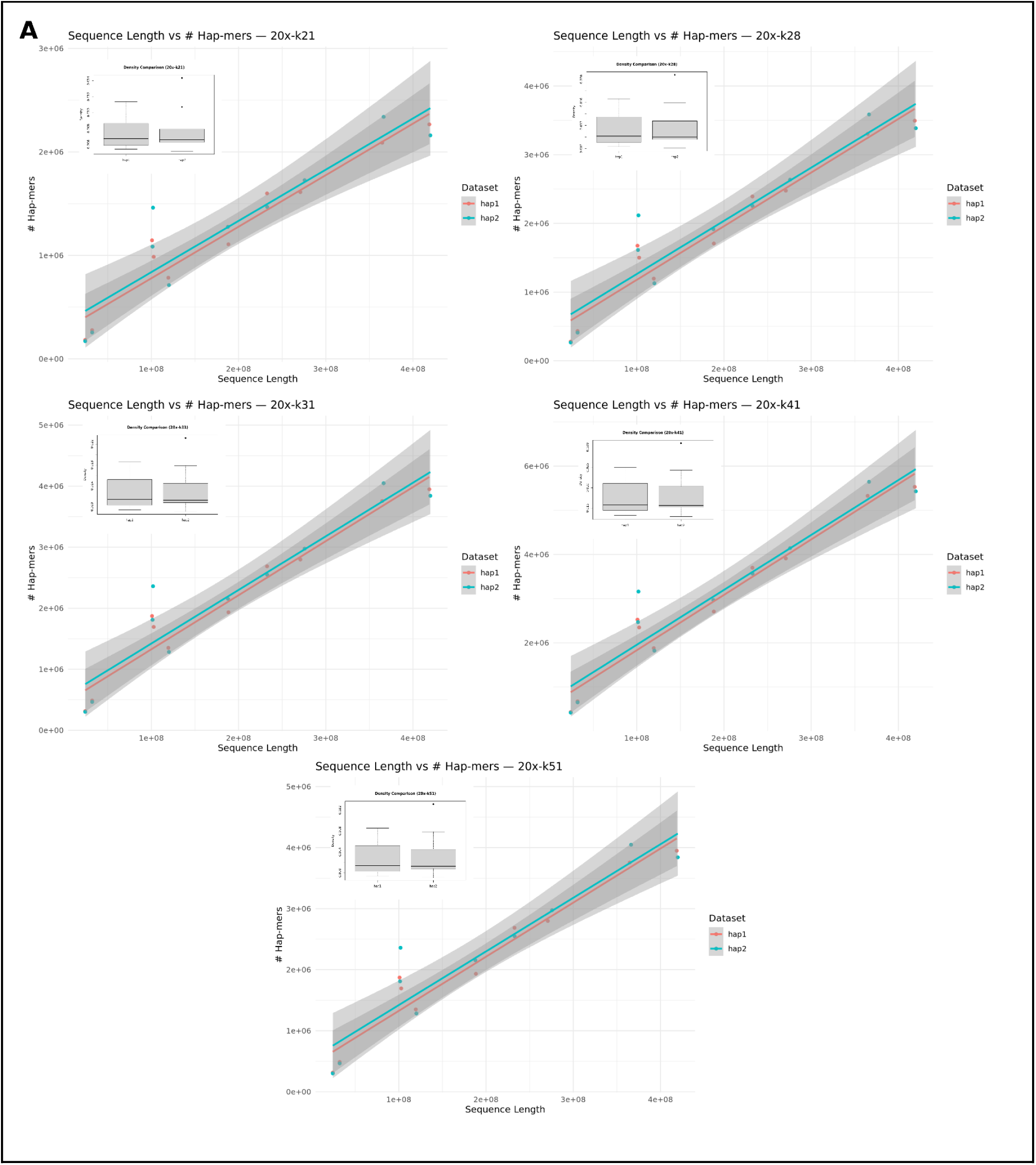
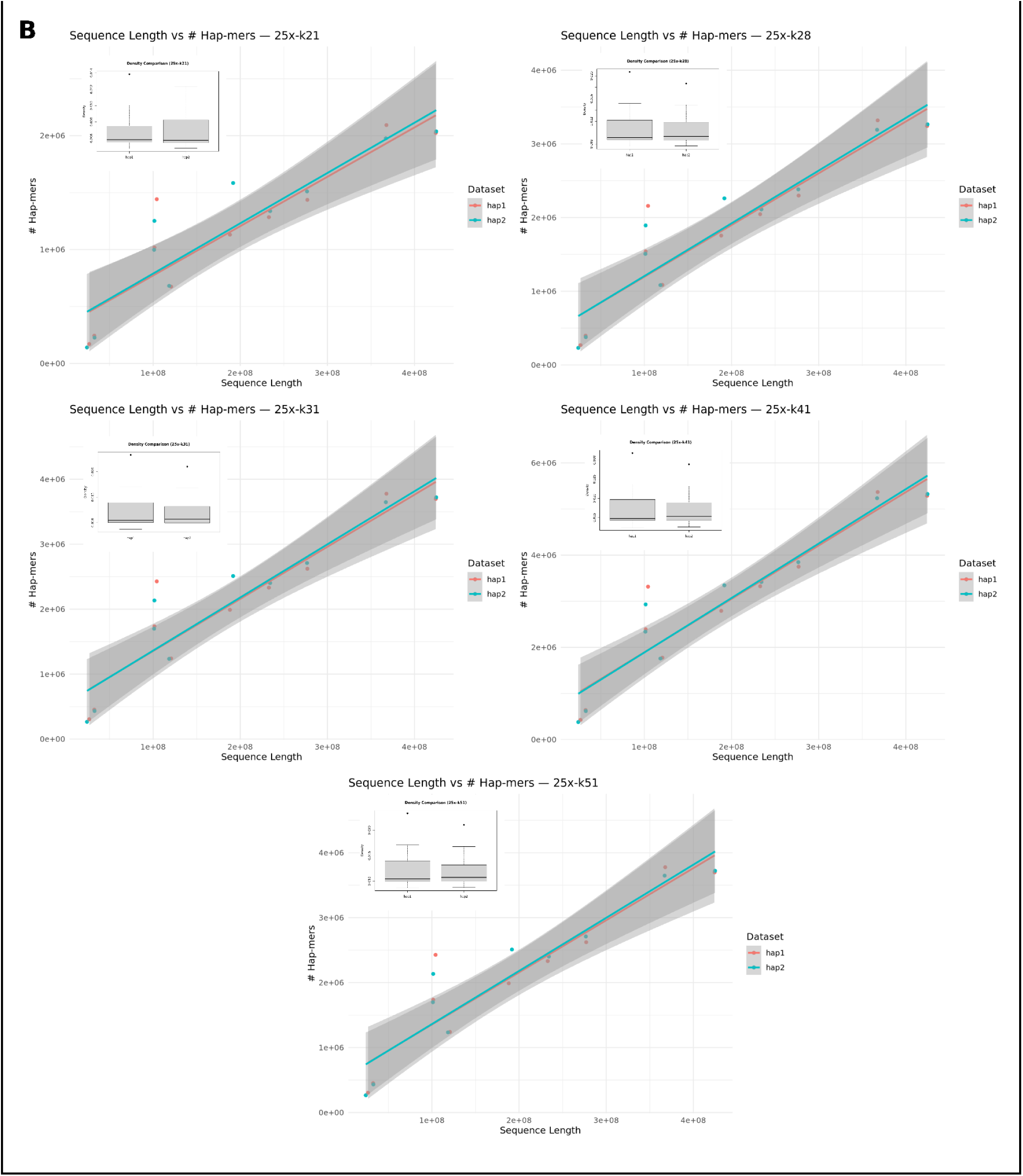
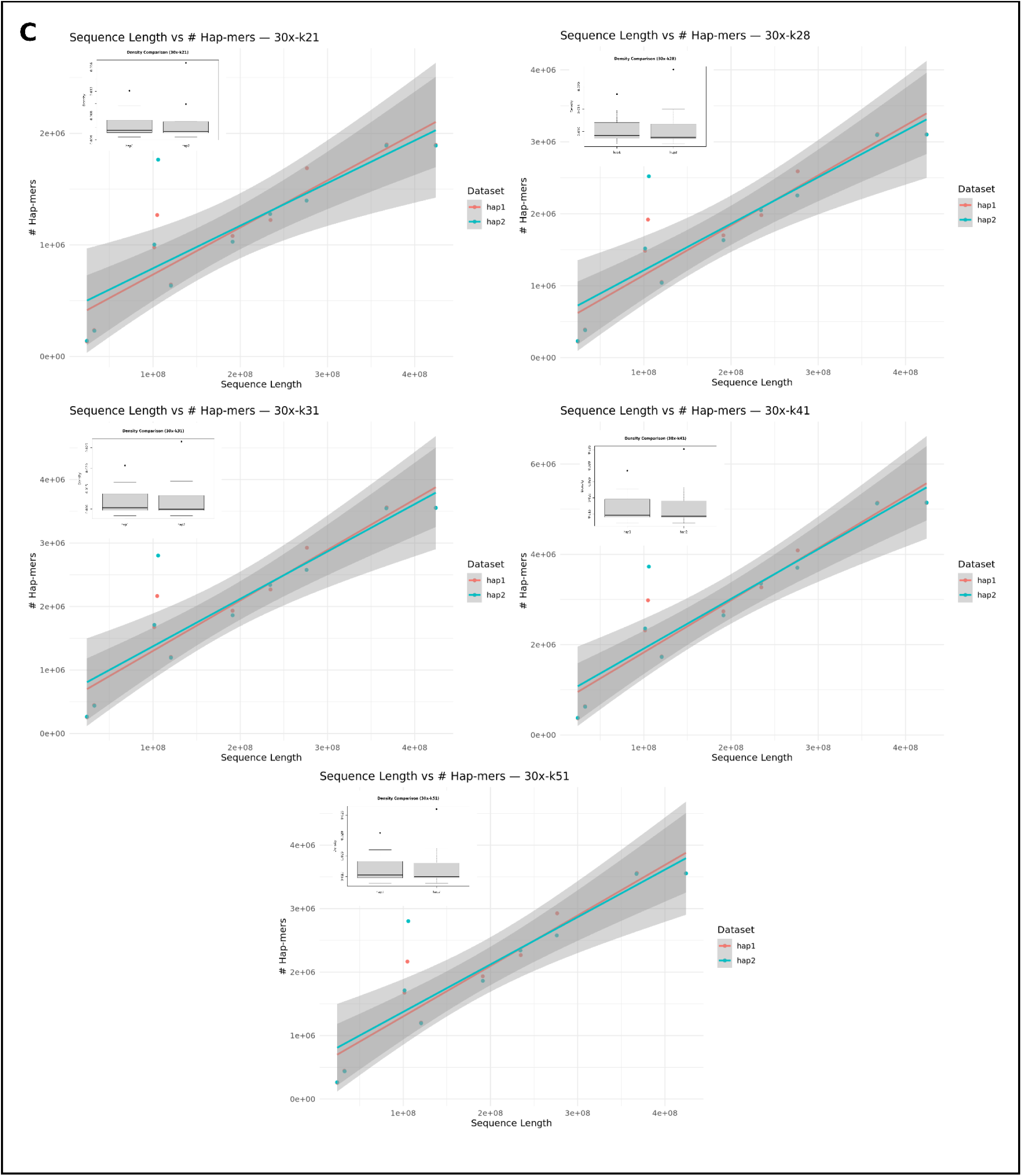
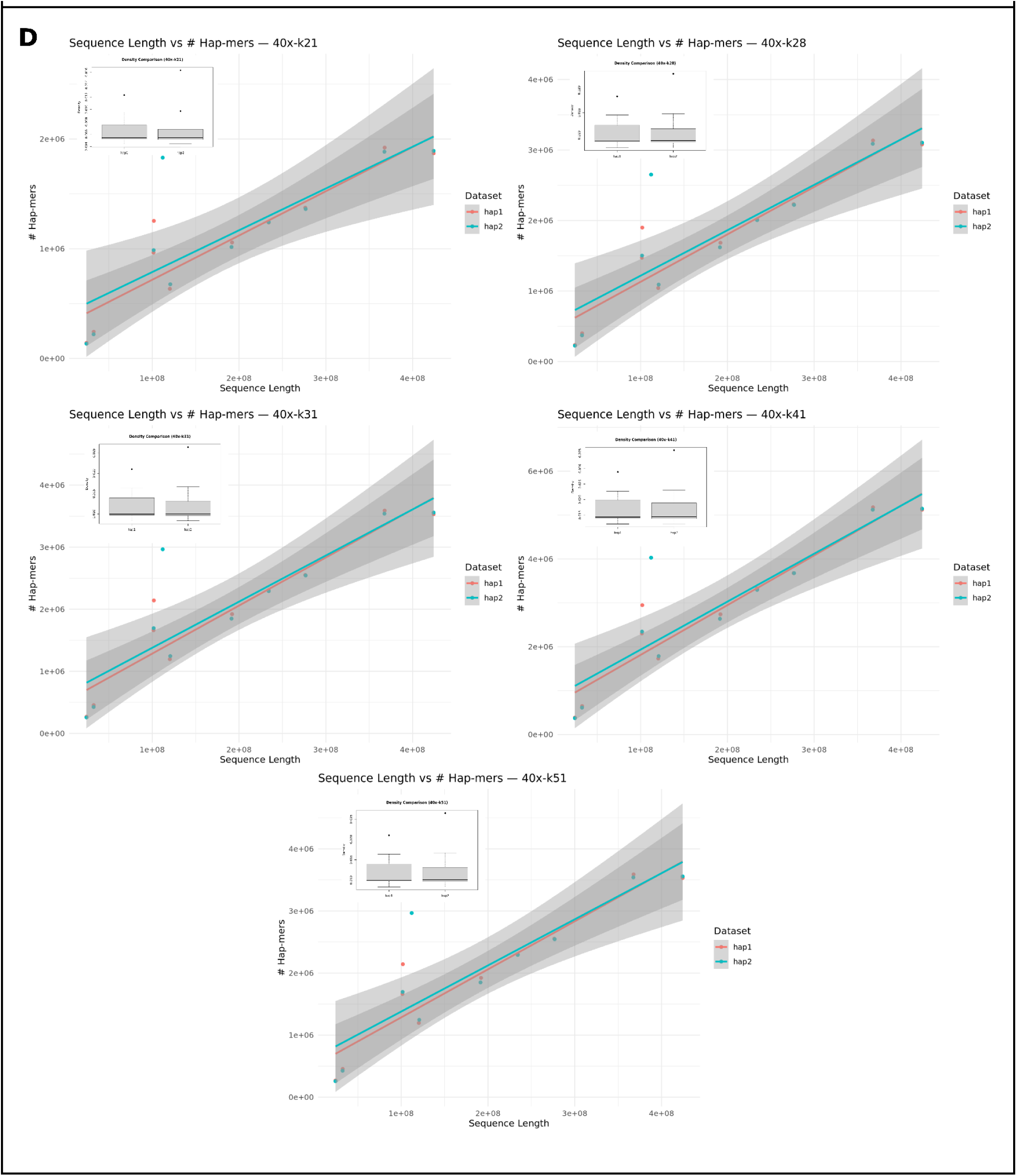
Co-exploration of the effects of varying average sequencing depth of the assembly and kmer lengths used by SCINKD in Anniella stebbinsi. (A) 20x coverage, (B) 25x coverage, (C) 30x coverage, (D) 40x coverage. At 20x coverage, it’s difficult to definitively observe signals from one of the two haplotypes at any kmer length, while at k21 at all sequencing depths, sex chromosome signal is obscured. The anticipated SCINKD signal remains robust at ≥25x and ≥k28.

**Supplemental Figure 3:**
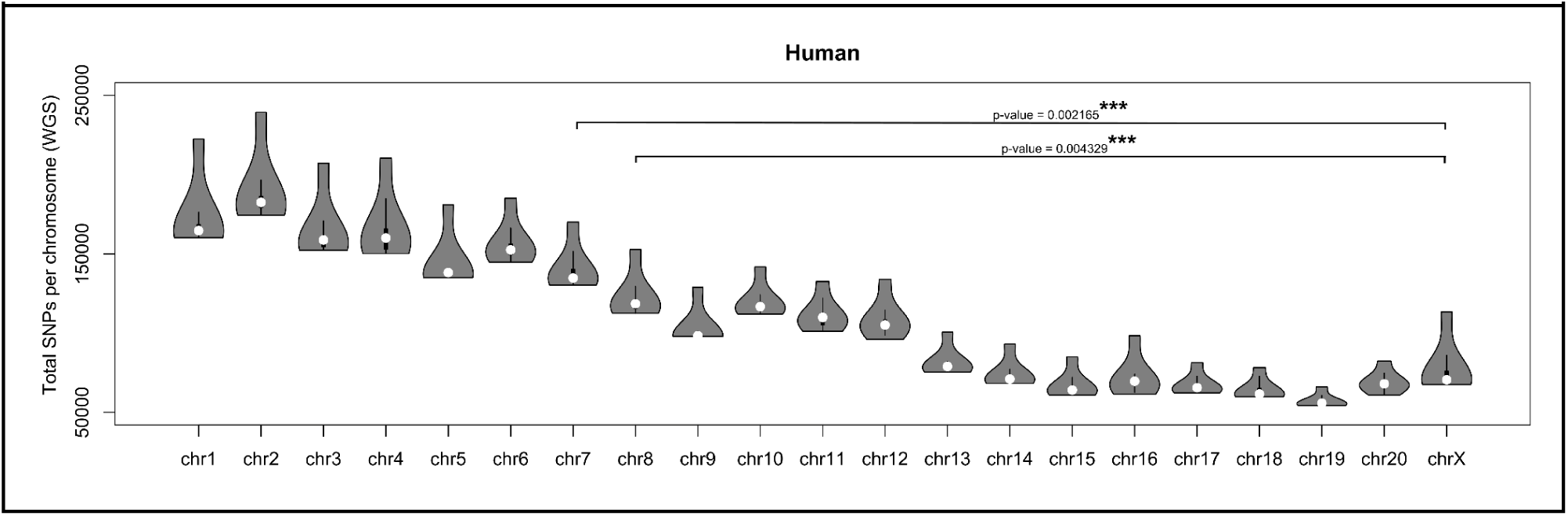
Preliminary data suggesting that the number of biallelic SNPs across individuals roughly correlates with chromosome size in humans using whole genome re-sequencing (WGS) in GTEx data. In addition, the X chromosome contains fewer SNPs than expected based on its length, which is the approximate length of chr7 in GRCh38.

**Supplemental Figure 4:**
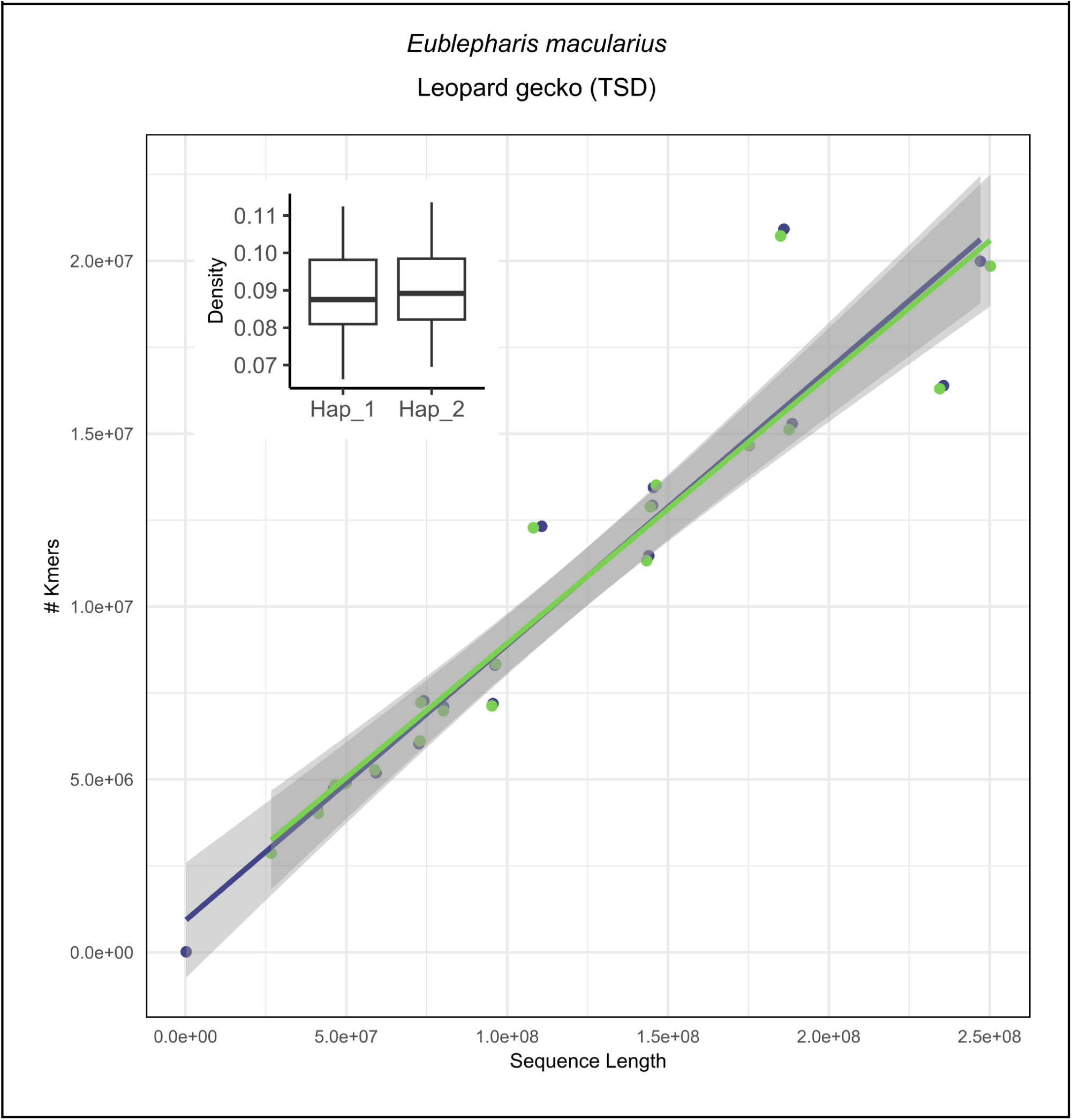
Confirmation of a strong correlation (R^2^ == 0.901) between hap-mers and chromosome length in a temperature-dependent species, the leopard gecko (*Eublepharis macularius*). Subtle variations between haplotypes are thought to be driven by poor phasing performance driven by low heterozygosity and/or sub-optimal coverage (30x) of PacBio HiFi data (Pinto et al. 2023).

**Supplemental Figure 5:**
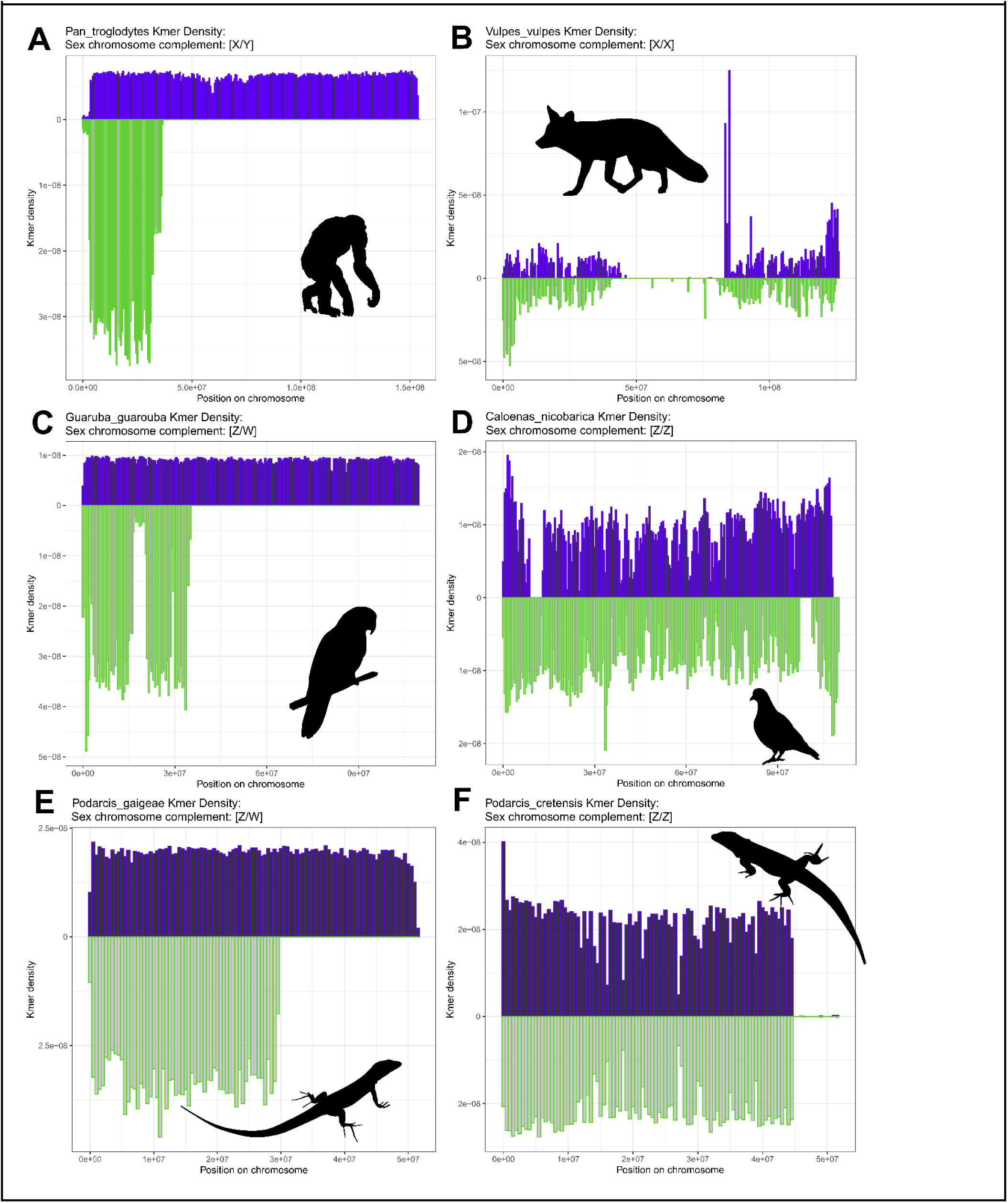
Mirror plots for both sexes of the heteromorphic systems detailed in Figure 2.

**Supplemental Figure 6:**
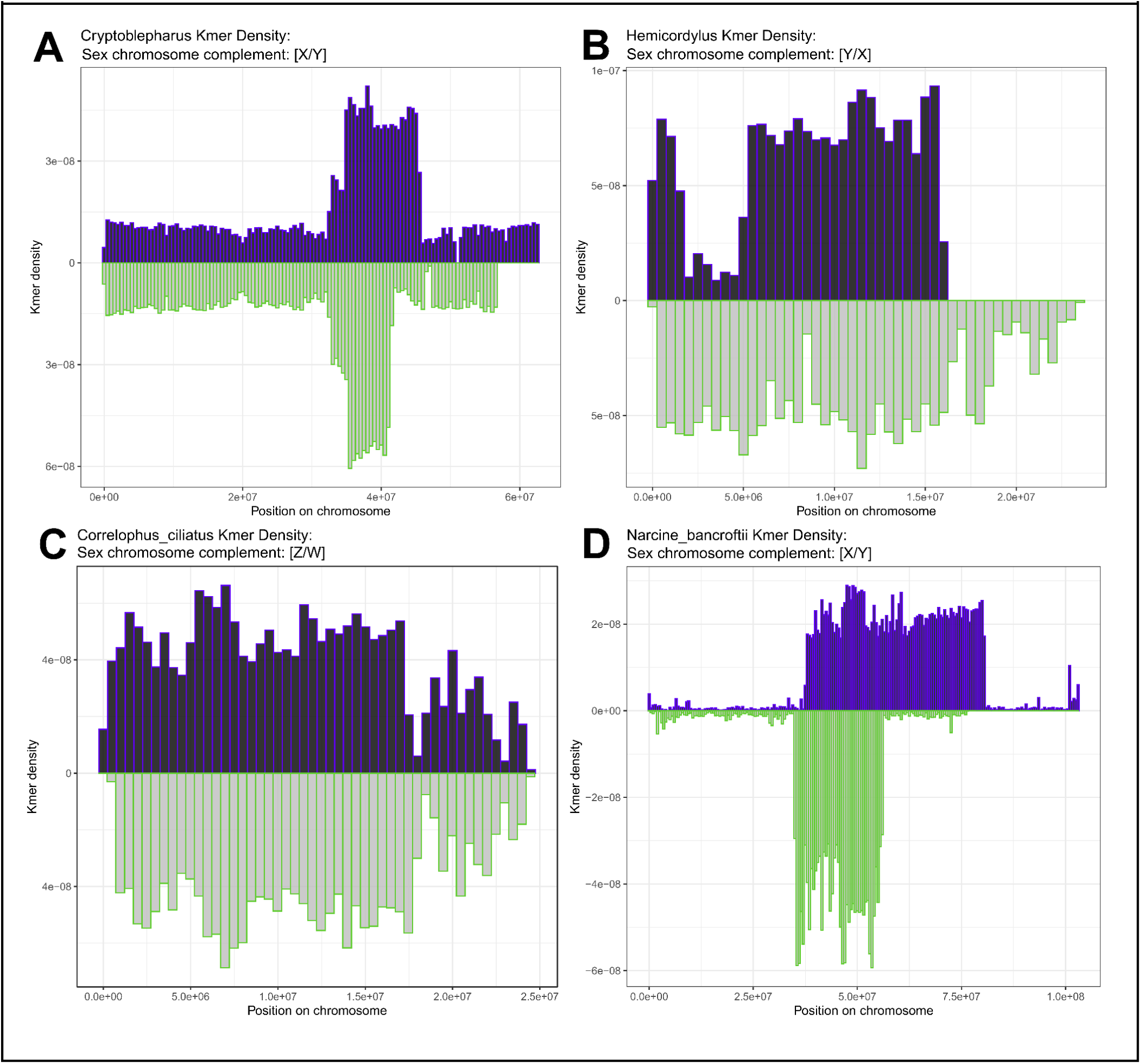
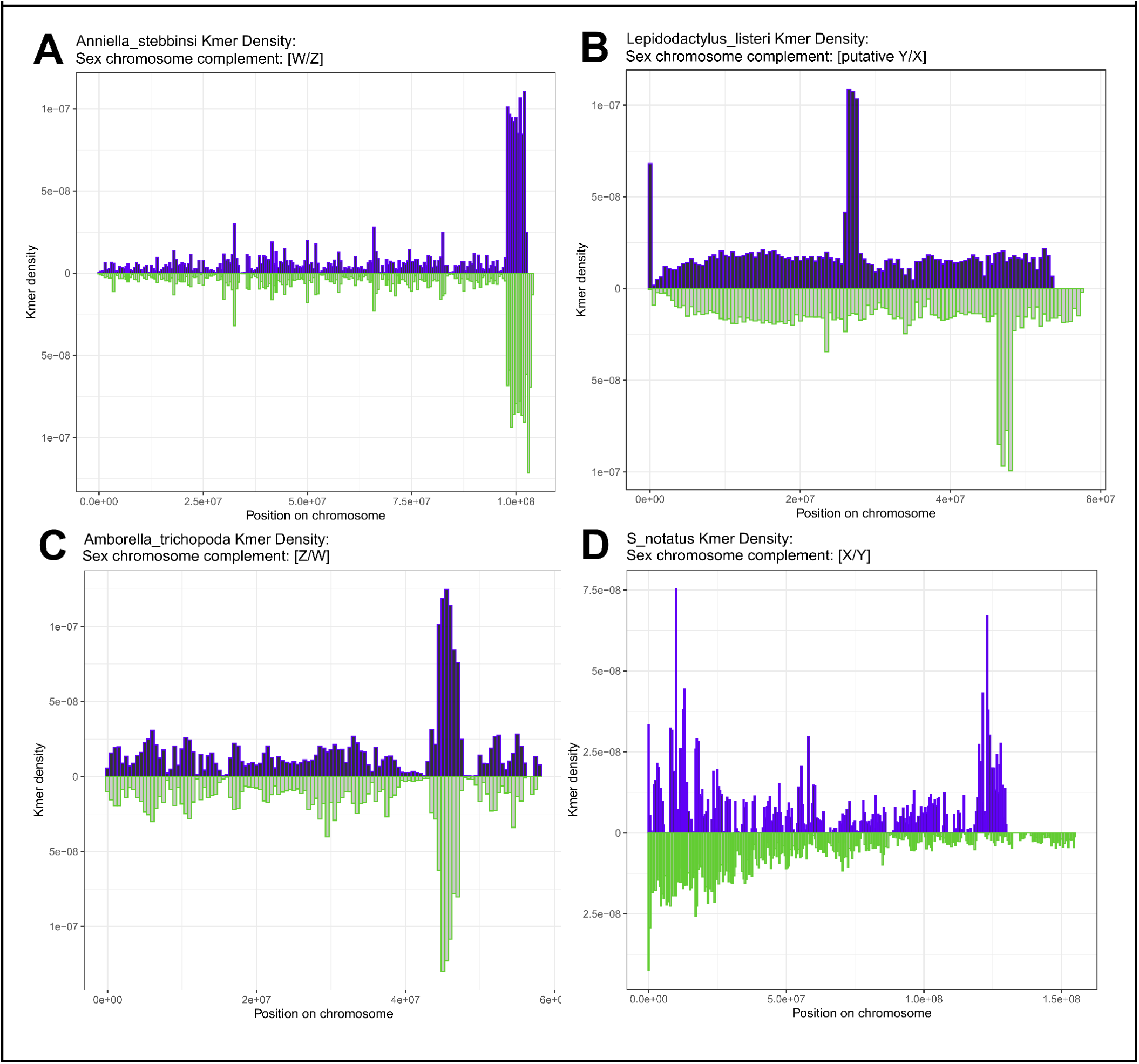
Single sex-only mirror plots.

